# FREEDA: an automated computational pipeline guides experimental testing of protein innovation by detecting positive selection

**DOI:** 10.1101/2023.02.27.530329

**Authors:** Damian Dudka, R. Brian Akins, Michael A. Lampson

## Abstract

Cell biologists typically focus on conserved regions of a protein, overlooking innovations that can shape its function over evolutionary time. Computational analyses can reveal potential innovations by detecting statistical signatures of positive selection that leads to rapid accumulation of beneficial mutations. However, these approaches are not easily accessible to non-specialists, limiting their use in cell biology. Here, we present an automated computational pipeline FREEDA (Finder of Rapidly Evolving Exons in De novo Assemblies) that provides a simple graphical user interface requiring only a gene name, integrates widely used molecular evolution tools to detect positive selection, and maps results onto protein structures predicted by AlphaFold. Applying FREEDA to >100 mouse centromere proteins, we find evidence of positive selection in intrinsically disordered regions of ancient domains, suggesting innovation of essential functions. As a proof-of-principle experiment, we show innovation in centromere binding of CENP-O. Overall, we provide an accessible computational tool to guide cell biology research and apply it to experimentally demonstrate functional innovation.

## Introduction

Purifying selection eliminates deleterious non-synonymous mutations, leading to conservation of amino acid sequence. In contrast, positive selection results in the accumulation of non-synonymous mutations that lead to functional innovation and adaptation (reviewed in (Nielsen et al., 2005)). Compelling examples of how positive selection regulates protein function come from studying host-pathogen genetic conflicts. In these evolutionary arms races, positive selection leads to rapid accumulation of mutations in both viral proteins that help infect the host and host proteins that help evade the infection (reviewed in (Daugherty and Malik, 2012; Sironi et al., 2015)). To experimentally test functional innovation, evolutionary biologists swap protein regions (or entire alleles) from closely related species that are highly diverged due to positive selection. This approach generates an “evolutionary mismatch” between the divergent protein and the cellular environment, revealing which protein function has undergone adaptation (reviewed in (Brand and Levine, 2021)). For example, swapping a region of the TRIM5 protein between human and rhesus monkey revealed that positive selection shapes its role in fighting species-specific retroviral infections (Sawyer et al., 2005; Stremlau et al., 2005; Yap et al., 2005). Remarkably, even single residues under positive selection can lead to adaptation, as in the human anti-viral MAVS protein that evolves to evade infection with hepaciviruses (Patel et al., 2012). These examples illustrate that innovation-guided functional analyses can complement more traditional conservation-guided approaches in revealing regulation of protein function.

Genetic conflicts, like those between host and pathogen, result in recurrently changing selection pressure and recurrent adaptation of proteins regulating essential cellular processes. For example, pressure to maintain genome integrity at fertilization fuels a sexual conflict between paternal proteins that adapt to maximize the chance of fertilizing the egg and maternal proteins that adapt to prevent entry of more than one sperm (reviewed in (Carlisle and Swanson, 2020)). Similarly, selfish genetic elements such as transposons constantly disrupt genome integrity, leading to intra-genomic conflicts and recurrent adaptation of DNA packaging proteins (reviewed in (Brand and Levine, 2021)). Centromere DNA sequences can also act as selfish elements, raising the possibility of intra-genomic conflict with centromere-associated proteins. Centromeres are repetitive DNA regions that direct chromosome segregation in mitosis and meiosis by assembling kinetochores, multi-protein structures that connect to spindle microtubules. Despite their essential function, centromeric DNA and proteins evolve rapidly across taxa, suggesting an evolutionary pressure to recurrently innovate. The centromere drive hypothesis proposes that selfish centromeric DNA sequences achieve non-Mendelian segregation during asymmetric female meiosis, increasing their transmission to the egg. Fitness costs imposed by this selfish behavior would lead to recurrent adaptation of centromeric proteins to suppress the costs (Henikoff et al., 2001). While there is experimental evidence for selfish centromeric DNA, the impact of positive selection on centromeric protein function remains largely untested (reviewed in (Dudka and Lampson, 2022)).

The scarcity of experimental studies of adaptive evolution in centromeric proteins, in contrast to our increasingly detailed understanding of their conserved functions (Kixmoeller et al., 2020; McKinley and Cheeseman, 2016; Mellone and Fachinetti, 2021), reflects the general focus of cell biology research on protein conservation rather than innovation. This discrepancy is due in part to challenges in designing experiments to demonstrate functional consequences of positive selection, but also to the complexity of methods needed to distinguish positive selection from neutral evolution of protein coding sequences (reviewed in (Anisimova and Liberles, 2012)). A widely used method calculates the rate ratio of non-synonymous (dN) to synonymous (dS) substitutions per codon (dN/dS ratio; (Goldman and Yang, 1994; Kimura, 1977; Muse and Gaut, 1994)) using multiple sequence alignment of homologous proteins from closely related species (orthologues). This approach assumes that synonymous mutations are neutral, while deleterious non-synonymous mutations are purged by purifying selection. Recurring non-synonymous substitutions within the alignment suggest recurrent adaptation to a constantly changing selection pressure (types of recurrent evolution are discussed in (Maeso et al., 2012)). Well-established computational suites such as PAML (Phylogenetic Analysis by Maximum Likelihood; (Yang, 2007)) and HyPhy (Hypothesis Testing using Phylogenies; (Pond et al., 2005)) offer a number of tools that can reliably detect positive selection but are seldom used by cell biologists because expertise in computational biology and molecular evolution is required to generate the input data, and the output is not provided in an intuitive visual format.

Automated molecular evolution pipelines that incorporate the abovementioned tools have been developed (see Supplementary Figure 1 for a non-exhaustive list), but their complexity and the need for user-provided input still renders them inaccessible to experimental cell biologists with limited computational skills. Increasing this access requires a “one-click” application that: 1) offers a simple graphical user interface, 2) fully automates input preparation, 3) finds orthologues despite the lack of genomic annotations, 4) reduces parameterization, and 5) provides intuitive visual representation of the output. Here, we present FREEDA (Finder of Rapidly Evolving Exons in De novo Assemblies), a fully automated, end-to-end pipeline designed for cell biologists seeking to apply an evolutionary lens by testing for evidence of positive selection in their favorite proteins. FREEDA provides the key functionalities listed above, including a unique ability to map residues under positive selection onto any predicted protein structure. As a proof-of-principle, we first use FREEDA to map positive selection across centromeric proteins in rodents, as mice are currently the only experimentally tractable cell biological model system for centromere drive (reviewed in (Dudka and Lampson, 2022)). Guided by these computational analyses, we use the evolutionary mismatch approach to provide experimental evidence of functional innovation in the centromeric protein CENP-O.

## Results

### Overview of the FREEDA pipeline

FREEDA is a stand-alone application with an intuitive graphical user interface (GUI) operating on UNIX systems (MacOS and Linux; Windows users please see documentation). An overview and documentation of the pipeline are provided here: https://ddudka9.github.io/freeda/ with a more detailed walkthrough in Methods. FREEDA first downloads the reference genome of the user-selected species and prepares input data for the gene of interest by connecting to genomic, protein, and protein structure databases (Fig. 1; blue). Next, FREEDA downloads a preselected set of non-annotated genome assemblies related to the reference species, performs a BLAST search for orthologues of the gene of interest, and uses the reference species data to find orthologous sequences (Fig. 1; orange). Combining several well-established molecular evolution tools, FREEDA aligns all coding sequences, builds phylogenetic trees, and determines whether any residues change recurrently throughout the phylogeny – a signal of positive selection (Fig. 1; brown). Key results are displayed within the GUI (Fig. 2A), and all results are saved into the “Results-current-date” folder generated in a location selected by the user (“Set directory”; Fig. 2A). These files include the BLAST output, nucleotide alignment, phylogenetic tree, protein alignment, residue mapping onto reference coding sequence, and residue mapping onto protein structure. The raw data and intermediate alignment files are saved in the “Raw_data” folder. Since FREEDA finds orthologues by downloading entire genomic assemblies, the user is advised to select an external data storage device (e.g., a hard drive) when setting the directory. Internet connection is also required to allow communication with various databases (Fig. 1).

**Figure 1.**
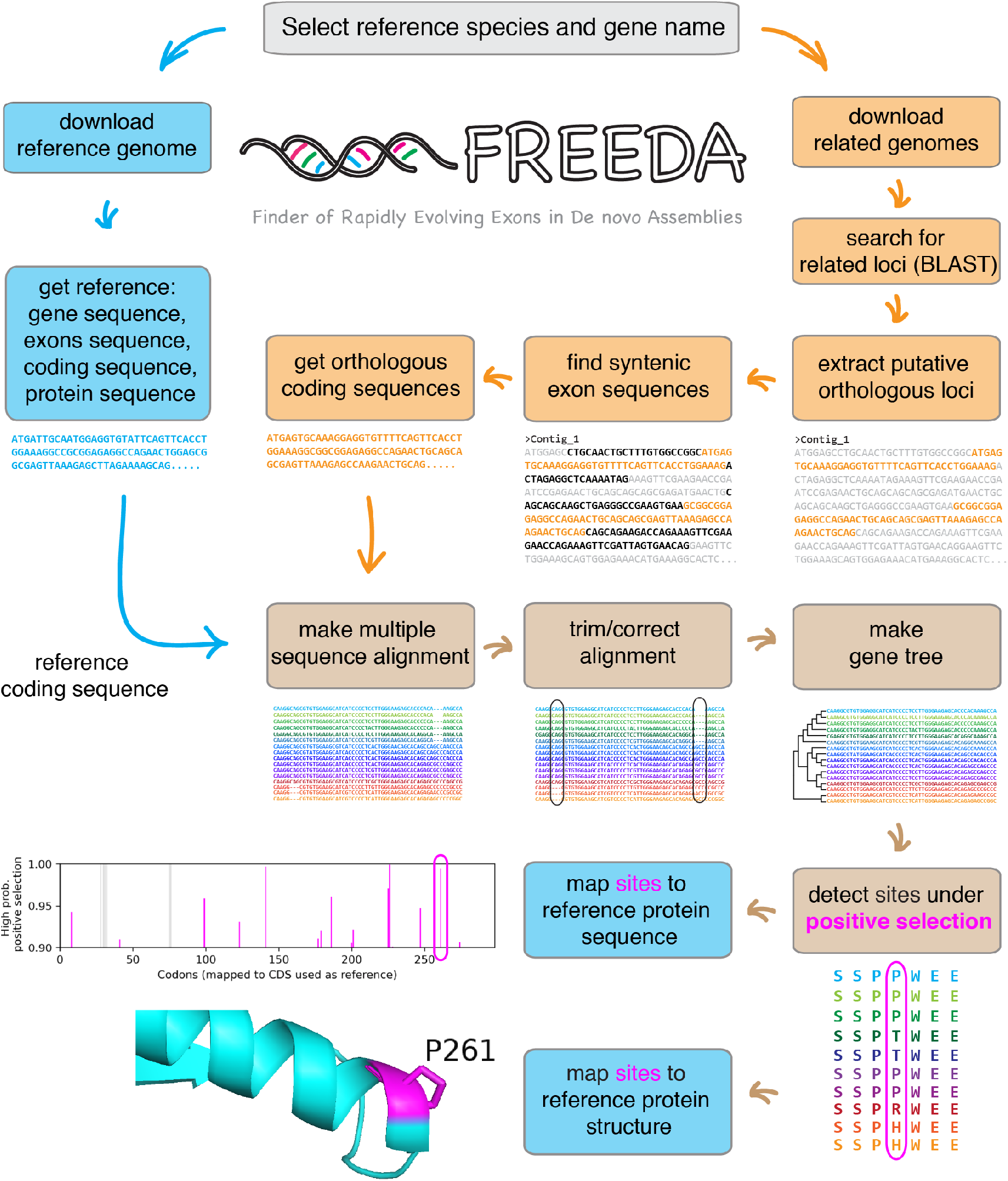
Overview of the FREEDA pipeline. The schematic shows the main steps, with more details in Methods. Launching the FREEDA application opens a graphical user interface (grey), which prompts selection of a reference species and a gene name. First, operating on the selected reference species (blue path), FREEDA downloads the genome and uses it to curate input for the gene of interest (protein sequence, exon sequences, coding sequence, and gene sequence). Second, operating on closely related species (orange path), FREEDA downloads non-annotated genomes, searches for putative orthologous loci, retrieves these loci, finds syntenic (matching the reference locus) exons, and assembles coding sequences of orthologous genes based on the intron-exon boundaries known for the reference gene. Third, operating on the multiple coding sequences (brown path), FREEDA makes and curates a multiple sequence alignment, generates a phylogenetic gene tree, and detects sites under positive selection using established models measuring the rate ratio of non-synonymous to synonymous substitutions. Fourth, FREEDA maps these sites onto both the reference coding sequence and the structure prediction of the reference protein.

**Figure 2.**
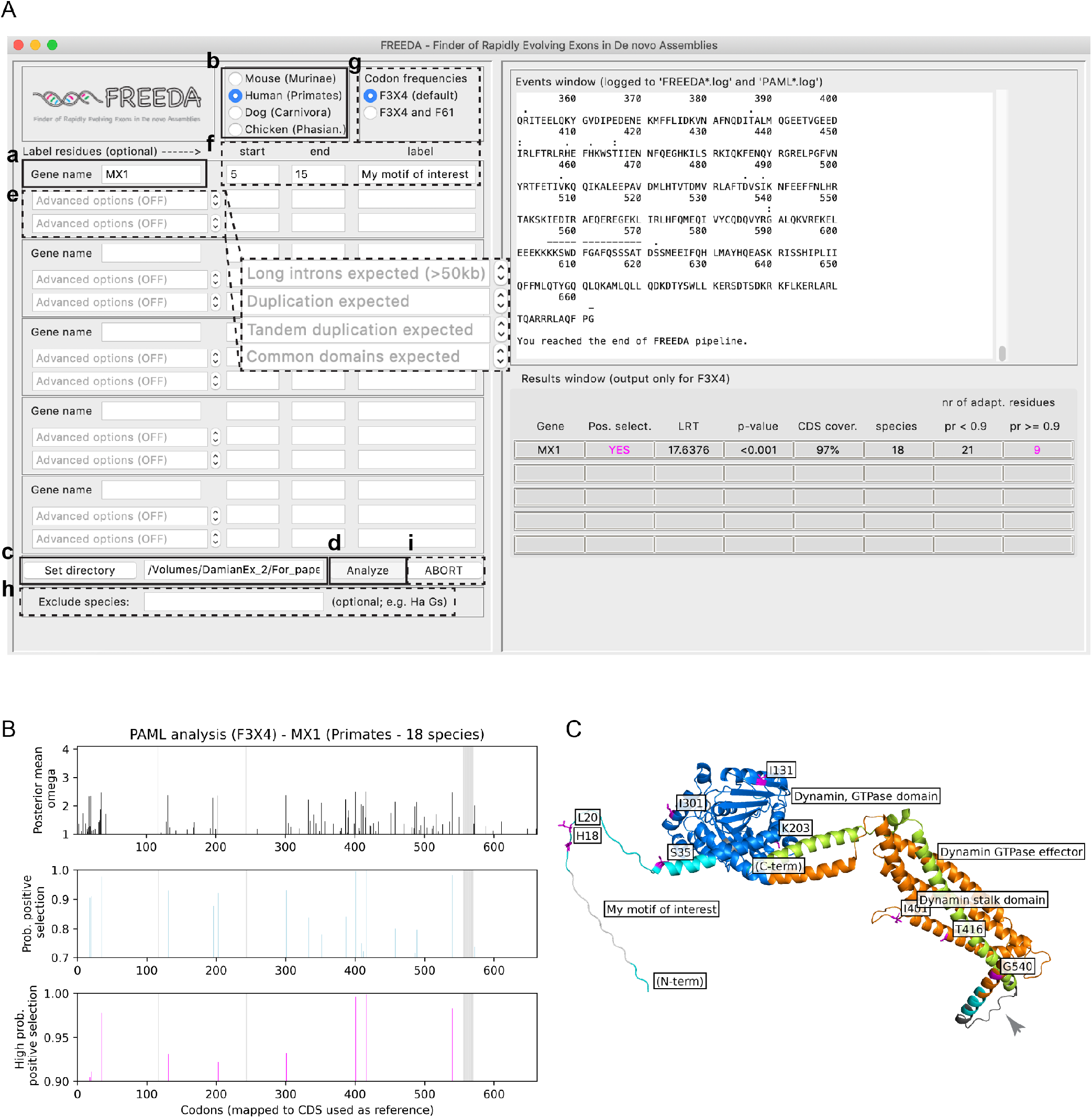
Example analysis of the primate *MX1* gene. **A)** FREEDA’s graphical user interface is divided into an input window (left half) and an output window (right half). The input window is used to provide a gene name (a), select the reference species (b), indicate where to save the data (c), and start the analysis (d). Optionally, the user can select advanced features (”Duplication expected”, “Tandem duplication expected”, “Long introns expected (>50kb)” and “Common domains expected”; see Documentation) (e), label up to 3 regions of choice on the protein structure (f), select an additional codon frequency model (g), exclude species from the analysis (h), and abort the analysis (i). The output window shows current tasks (“Events window”, top) and key results for each gene (“Results window”, bottom). **B**) Putative adaptive sites are mapped onto the reference coding sequence. Graphs show recurrently changing residues (top, black bars), residues that are likely to have evolved under positive selection (middle, blue bars, probability ≥ 0.7), and most likely targets of positive selection (bottom, magenta bars, probability ≥ 0.9). Grey bars in all graphs show residues removed from the analysis. **C**) Residues with the highest probability of positive selection (magenta) are mapped on the structural prediction model of the MxA protein (encoded by *MX1*) from AlphaFold. The N- and C-termini and known domains are automatically annotated, in addition to the user-specified region (“My motif of interest”). Regions removed from the analysis (arrowhead) are colored dark grey. To clearly show residues under positive selection, labels were modified manually in PyMOL (raw output is shown in Fig. S2E).

### Advantages over existing automated pipelines

Several features distinguish FREEDA from currently available automated pipelines (Supplementary Figure 1). First, FREEDA is fully automated, requiring only a gene name, and distributed as a self-contained application that does not require installation or compilation of any additional programs, except for a straightforward installation of the widely used protein structure viewer PyMOL (The PyMOL Molecular Graphics System, Version 2.0 Schrödinger, LLC) for MacOS users. Second, FREEDA uses a defined set of non-annotated genomic assemblies that ensures high statistical power of the analysis while absolving the users from manually curating their input. As new genomic assemblies become available, they will be incorporated into new FREEDA releases. Third, FREEDA automatically maps residues under positive selection onto protein structure models by querying the AlphaFold database containing structure predictions for nearly all mouse and human proteins (Jumper et al., 2021). Finally, by providing a simple GUI, FREEDA minimizes complexity compared to currently available pipelines, while offering a restrained number of advanced options (Fig. 2A). Therefore, the user may consider FREEDA as an entry point to performing the first molecular evolution analyses of their proteins of interest.

### FREEDA validation using genes with previously defined evolutionary histories

To demonstrate that FREEDA’s simplicity does not compromise its functionality, we used a dataset of 23 primate (*Simiiformes*) genes whose signatures of positive selection (or lack thereof) have been previously defined. The dataset included 19 genes curated to validate the DGINN pipeline (Detect Genetic INNovations; (Picard et al., 2020)). Analyzing a set of 19 primate species, FREEDA found 18 orthologues with 98% sequence coverage (median values; Supplementary Table 1). Consistent with the literature, FREEDA found evidence of positive selection in *TRIM5*, *MAVS*, *SAMHD1*, *IFI16*, *ZC3HAV1*, *RSAD2*, GBP5, *MX1*, *APOBEC3F* and *NBN* (Supplementary Table 1). Although previous studies also reported positive selection in *BST2* using 9 primate species (Gupta et al., 2009; van der Lee et al., 2017), FREEDA only found a weak signature of positive selection (p=0.0864; Supplementary Table 1), possibly due to positive selection operating mostly in the lineage of New World Monkeys in this gene (Liu et al., 2010). Of six genes whose evolutionary history is less clear, with results dependent on the method used (Picard et al., 2020), FREEDA found evidence of positive selection in only one (*SERINC3*; Supplementary Table 1), highlighting the stringency of the analysis. Of six genes that have been shown to not evolve adaptively, FREEDA detected positive selection in one: *TREX1*, a nuclease that guards genome integrity (Picard et al., 2020); Supplementary Table 1). One of the most likely adaptive residues (S166; probability = 0.97; Supplementary Table 1) is proximal to a primate-specific DNA binding site (R164; (Zhou et al., 2022)), suggesting adaptive evolution of DNA recognition. Consistent with our finding, divergent DNA binding sites in *TREX1* regulate DNA recognition (Zhou et al., 2022). We suspect that disparity with previous *TREX1* analyses is due to differences in the alignment algorithms used (see Methods). Altogether, these analyses validate our pipeline using an objectively selected dataset and provide additional insight into the evolutionary history of these genes.

Rigorous identification of orthologues, as opposed to paralogues formed by a duplication event, is critical to avoid false signatures of positive selection. FREEDA demonstrated the ability to resolve ancient duplications by correctly distinguishing *HERC5* orthologues from *HERC6* paralogues present within our test dataset ((Picard et al., 2020); Additional Supplementary Materials). Additionally, we tested if FREEDA could resolve tandem duplications (duplicated genes located side by side) and retro-duplications (intron-less mRNA that was reverse-transcribed and inserted back into the genome). Using the “Tandem duplication expected” option (see GUI; Fig. 2A), FREEDA successfully distinguished primate genes *H4C1* and *H4C2*, both encoding histone H4 and located merely 5kb apart with 85% nucleotide sequence identity (Additional Supplementary Materials). Visual examination of the nucleotide alignment revealed that one *H4C2* orthologue lost its start codon and likely pseudogenized. In such cases, the user may choose to re-run the analysis using the “Exclude species” option (see GUI; Fig.2A). We further used the “Duplication expected” option (see GUI; Fig.2A) to show that FREEDA could correctly distinguish *KIF4A*, encoding a kinesin motor, from its retro-duplicate *KIF4B*. These genes are an example of a recent duplication that occurred in a common ancestor of primates, retaining 96% nucleotide sequence identity ((Florio et al., 2018); Additional Supplementary Materials). We also found that *KIF4B*, and not *KIF4A*, evolves under positive selection, suggesting adaptive evolution spurred by a duplication event. While human KIF4A kinesin regulates chromosome segregation (Mazumdar et al., 2004), cellular transport (Peretti et al., 2000) and anti-viral response (Gad et al., 2022), KIF4B remains poorly studied. Finally, analyzing our test dataset showed FREEDA’s sensitivity to gene losses, as no complete coding sequence was retrieved for *GBP5* in Old World Monkeys, which was previously shown to be lost in this branch ((Picard et al., 2020); Supplementary Table 1). Together, these analyses demonstrate that FREEDA reliably finds orthologous sequences.

To further validate accuracy of the pipeline at the level of single residues, we compared specific sites under positive selection found by FREEDA to those previously mapped in *MAVS*, *MX1*, *SAMHD1* and *TRIM5* (Laguette et al., 2012; Lim et al., 2012; Mitchell et al., 2012; Patel et al., 2012; Sawyer et al., 2005; van der Lee et al., 2017). Exact matching of probabilities for each residue was not expected due to differences in algorithms for aligning orthologous sequences (see Methods for details). Nevertheless, FREEDA found evidence of positive selection in all published sites, except for those located in regions removed from the alignment to ensure its high quality (5 residues in each *SAMHD1* and *MX1*; 2 residues in *TRIM5*; Fig. S2A-E; Supplementary Table 2). Using *MX1* as an example (Fig. 2A), FREEDA maps detected sites under positive selection onto the reference coding sequence (Fig. 2B) and onto structural prediction models generated by AlphaFold (Fig. 2C; (Jumper et al., 2021)). Overall, in these analyses we demonstrate that FREEDA can retrieve expected sites under positive selection, and we showcase FREEDA’s key results visualization features.

Finally, we tested if FREEDA can reliably detect positive selection in mouse proteins using mouse-related genomes (*Murinae*). As a test dataset, we selected 104 centromeric genes, 42 of which have been previously analyzed using a smaller number of species (up to 11; (Kumon et al., 2021)). Analyzing a set of 19 *Murinae* species, FREEDA found 16 orthologues and 94% sequence coverage (median values). Consistent with our previous findings of pervasive evolutionary innovation across the rodent centromere (Kumon et al., 2021), FREEDA found evidence of positive selection in 36/104 genes (Fig. 3; Supplementary Table 3). Corroborating our previous results, FREEDA detected positive selection in genes encoding CENP-C, CENP-I, CENP-T, HJURP, INCENP, MIS18BP1, KNL1 and SGO2. In contrast, DSN1 and NDC80 did not show signatures of positive selection. This discrepancy likely reflects a higher statistical power, due to more orthologues (up to 19) included in the analysis, to distinguish between positive selection and relaxation of purifying selection, both of which can result in sequence divergence. This high statistical power revealed several previously unknown targets of positive selection, including components of the fibrous corona, which helps capture microtubules (CENP-F, SPINDLY, ZWILCH, ROD, NUP85, NUP98 and ELYS; reviewed in (Kops and Gassmann, 2020)), microtubule motors (CENP-E, KIF2B and KIF18A) and protein kinases (AURKC and HASPIN). To further validate our findings, we repeated the analyses with rat as reference species. Since the quality of the available rat genome annotation is lower than that of mouse, FREEDA was able to collect reliable input data for only 89/104 genes. As expected, we found positive selection or lack thereof in almost exactly the same genes as when using mouse as reference (85/89 genes; see Discussion; Supplementary Table 3). Overall, these tests show that despite its simplicity for the user, FREEDA is a fully functional and dependable tool to detect positive selection.

**Figure 3.**
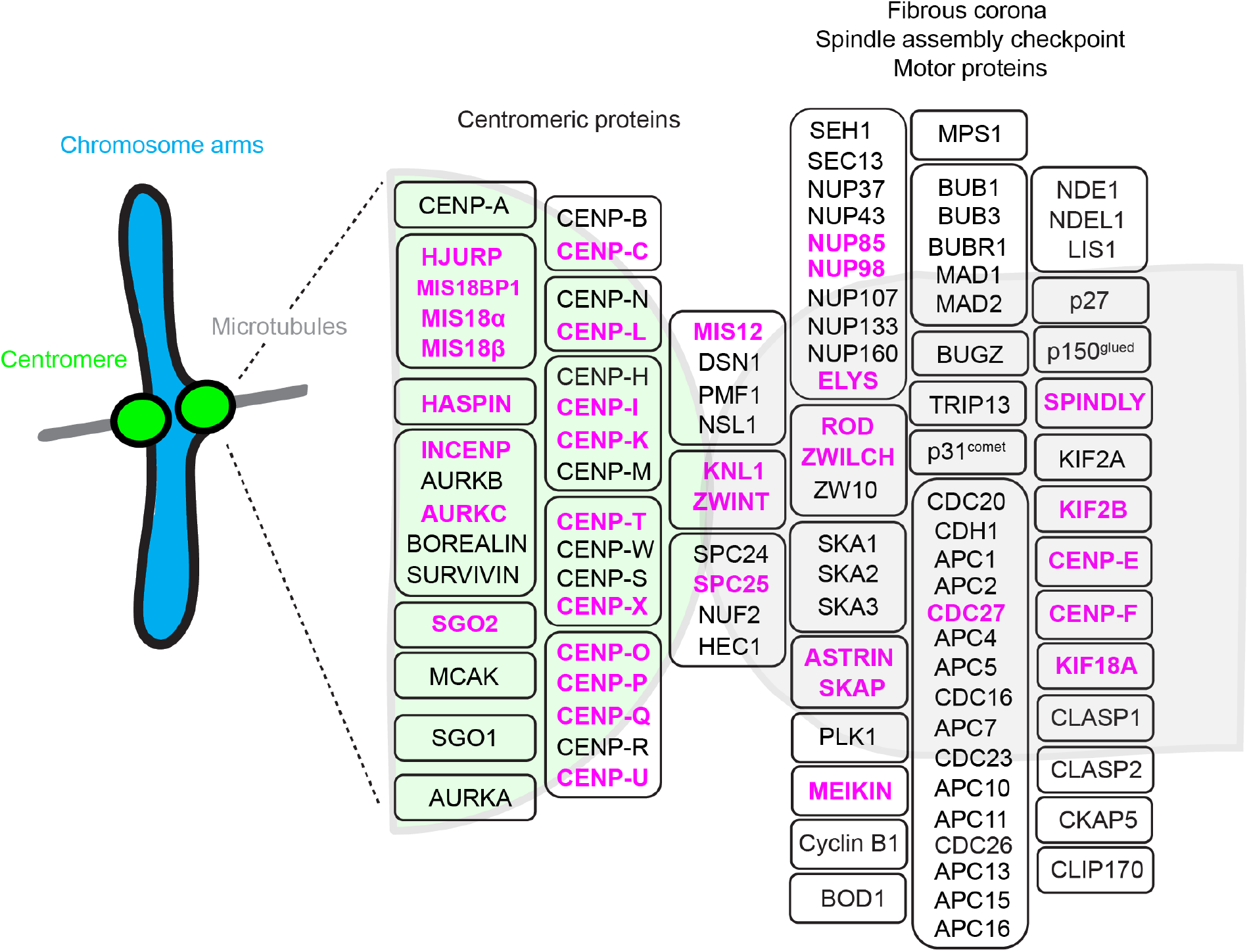
Positive selection across the rodent centromere. Centromere-associated proteins are grouped to reflect their approximate positions between chromatin and microtubules. Relative protein placement is informed by binding partners, but not all known interactions are depicted. Proteins forming complexes are grouped together. Magenta: proteins evolving under positive selection.

### Using FREEDA to derive evolution-guided hypotheses

To test if FREEDA can help derive evolution-guided hypotheses, we leveraged its ability to map residues under positive selection onto protein structures. We found evidence of positive selection within ancient (retained across long evolutionary timescales) protein domains of centromeric proteins, suggesting adaptive evolution of essential protein functions (Fig. 4A-F). For instance, we detected positive selection in the ancient Yippee domain (Roxstrom-Lindquist and Faye, 2001) of MIS18β (encoded by *Oip5*), which participates in centromere chromatin assembly (reviewed in (Zasadzinska and Foltz, 2017)). In addition to its divergent N- and C-termini, one of the most likely adaptive residues (R76; probability = 0.98) is located within the intrinsically disordered CXXC motif of the Yippee domain (Fig. 4A and B), which is required for MIS18 complex assembly at centromeres (Fujita et al., 2007; Stellfox et al., 2016; Subramanian et al., 2016).

**Figure 4.**
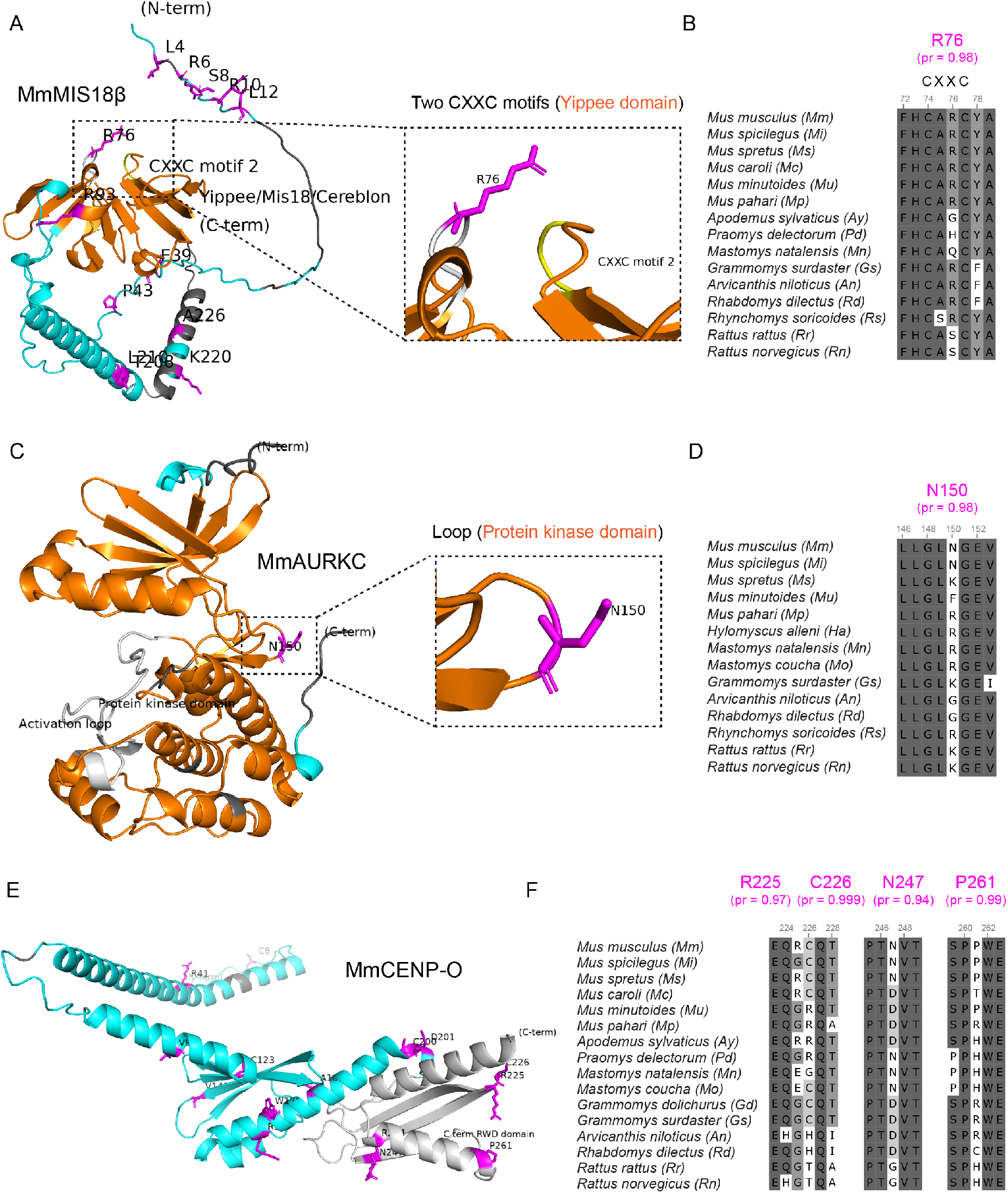
Positive selection in intrinsically disordered regions within ancient protein domains. Ribbon diagrams (**A, C, E**) show annotated structural prediction models of mouse proteins, generated automatically by FREEDA and visualized in PyMOL without manual modifications. Residues with high probability of positive selection are colored magenta, and a subset of these are shown in snippets of the multiple sequence alignments in *Murinae* (**B, D, F**). Dark grey: highly conserved residues; grey: less conserved residues; white: non-synonymous substitutions. **A** and **B**) MIS18β (MmMIS18β) shows the Yippee domain (orange) and two CXXC motifs (indicated by the user within the GUI, grey and yellow). The label for CXXC motif 1 is not visible to accommodate labeling of the R76 residue. The two CXXC motifs are enlarged with the R76 residue (magenta) within motif 1. **C** and **D**) AURKC (MmAURKC) shows the protein kinase domain (orange) and activation loop (indicated by the user within the GUI, grey). A loop within the protein kinase domain is enlarged, with N150 shown in the multiple sequence alignment. **E** and **F**) CENP-O (MmCENP-O) shows the C-terminal RWD domain (indicated by the user within the GUI, grey). The most likely adaptive residues of C-terminal RWD domain are shown in the multiple sequence alignment.

Similarly, we found strong evidence of positive selection in an intrinsically disordered loop of an ancient protein kinase domain (reviewed in (Taylor and Kornev, 2011)) in the meiosis-specific Aurora kinase C (AURKC, N150, probability = 0.98; Fig. 4C and D), which helps correct erroneous kinetochore-microtubule attachments (Balboula and Schindler, 2014). In contrast, we found no recurrent changes in the related AURKA and AURKB kinases (Fig. S3A-C). These data suggest that positive selection uniquely tunes the kinase activity of the specialized meiotic Aurora kinase, consistent with frequent adaptive evolution of reproduction genes (Jagadeeshan and Singh, 2005; Nielsen et al., 2005; Swanson and Vacquier, 2002).

Finally, we found evidence of positive selection within the ancient double RWD domain (RING-WD-DEAD; (Tromer et al., 2019)) of CENP-O, which regulates kinetochore-microtubule attachments by forming the CENP-OPQUR complex ((Amaro et al., 2010; Bancroft et al., 2015; Chen et al., 2021; Hori et al., 2008; Singh et al., 2021); Fig. 4E and F). RWD domains are prevalent structural modules that facilitate protein-protein interactions across the centromere (Schmitzberger and Harrison, 2012; Tromer et al., 2019). CENP-O shares a high structural similarity with its binding partner CENP-P, which also shows signatures of positive selection within its double RWD domain (Fig. S4A and B). Furthermore, some of the most likely adaptive residues flank highly structured α-helices and β-sheets in CENP-O and -P C-terminal RWD domains (Fig. 4E; Fig. S4A). Based on these results, we propose that positive selection regulates essential functions of centromeric proteins by acting on intrinsically disordered regions (such as loops and β-turns) within ancient domains, consistent with previous reports of frequent innovation of flexible regions of other proteins (Afanasyeva et al., 2018; Nilsson et al., 2011; Ridout et al., 2010). Altogether, we demonstrate that FREEDA can help derive evolution-guided hypotheses by highlighting protein domains with signatures of adaptive evolution.

### Using FREEDA to infer molecular mechanisms regulated by positive selection

Each of the proteins discussed above (MIS18β, AURKC and CENP-OP) functions as part of a complex. To infer mechanisms regulated by positive selection in this context, we aligned FREEDA-annotated protein structure predictions of mouse proteins (Fig. 4) to experimentally solved structures of their orthologues in complex with binding partners (see Methods for details). Two intrinsically disordered CXXC motifs within the Yippee domain of MIS18β together form a tetrahedral module whose four conserved cysteins bind a zinc ion (Subramanian et al., 2016), likely stabilizing protein conformation (Nguyen et al., 2020). Aligning mouse MIS18β to the crystal structure of the fission yeast MIS18 Yippee-like domain ((Subramanian et al., 2016); Fig. 5A and shows that the side chain of the positively selected R76 likely faces the opening of the tetrahedral module. This finding is consistent with XX residues regulating the function of CXXC motifs in other proteins (Quan et al., 2007). Alternatively, R76 could mediate MIS18α and MIS18β heterodimerization (Subramanian et al., 2016). Therefore, we hypothesize that positive selection favors amino-acid changes within the intrinsically disordered CXXC motif to modulate MIS18 complex stability. We also found recurrently changing, albeit less likely adaptive residues within the second CXXC motif of MIS18β (G135; probability = 0.88; Fig. 5A and B) and in the first CXXC motif of its binding partner MIS18α (S57; probability = 0.77; Fig. S5A-C), consistent with functional innovation of CXXC motifs. These data suggest that positive selection in the intrinsically disordered regions of the ancient Yippee domains regulates centromere assembly by modulating stability of the MIS18 complex.

**Figure 5.**
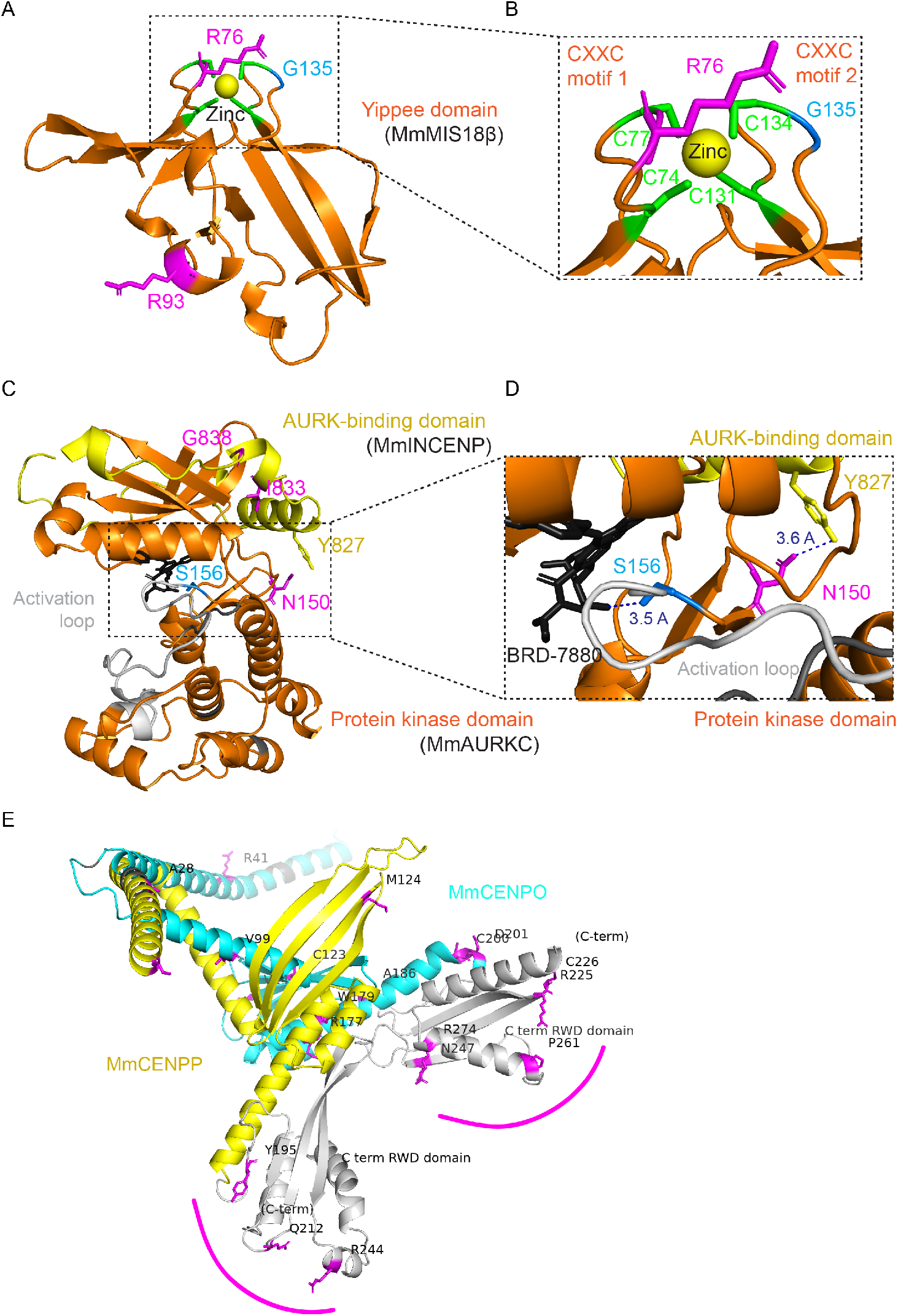
Putative molecular interactions of positively selected residues. **A**) Yippee domain (orange) of MmMIS18β aligned to the Yippee-like domain of fission yeast MIS18 (5JH0; (Subramanian et al., 2016)). FREEDA automatically annotates residues with high probability of positive selection (probability ≥ 0.9, magenta). Manual annotations show residues with lower probability of positive selection (probability ≥ 0.7, blue) and conserved cysteins (green). **B**) Enlarged CXXC motifs forming a tetrahedral module holding a zinc ion. **C**) MmAURKC protein kinase domain (orange, with activation loop in grey) and AURK-binding domain of mouse INCENP (MmINCENP; yellow), both aligned to the human AURKC-INCENP complex with the BRD-7880 inhibitor bound to the ATP-binding site (black; 6GR8; (Abdul Azeez et al., 2019)). Annotations show residues under positive selection (magenta, probability ≥ 0.9; blue, probability ≥ 0.7) and conserved residue Y827 of MmINCENP (yellow). **D**) Enlarged ATP-binding site. Dashed lines show closest distance from side chains of residues under positive selection (S156 or N150) to the BRD-7880 inhibitor or to Y827 of MmINCENP. **E**) MmCENP-O (blue) and MmCENP-P (yellow) aligned to human CENP-O and -P (HsCENP-O and -P) from the human CENP-OPQUR complex (7PB8; (Yatskevich et al., 2022)). Grey: C-terminal RWD domains; magenta: residues under positive selection (probability ≥ 0.9). Magenta arcs highlight residues under positive selection within the C-terminal RWD domains facing opposite sides of the heterodimer, which likely interface with other centromeric proteins.

AURKB and AURKC kinase activity requires binding to a conserved domain of INCENP (INner-CENtromere Protein; reviewed in (Krenn and Musacchio, 2015)). Aligning the MmAURKC protein kinase domain and MmINCENP AURK-binding domain to the crystal structure of the orthologous human domains (Abdul Azeez et al., 2019) shows the positively selected N150 side chain in close proximity to the conserved Y827 residue in MmINCENP. This finding suggests modulation of INCENP binding and therefore kinase activity by positive selection (Fig. 5C and D). The rodent AURKC activation loop also contains a recurrently changing, albeit less likely adaptive residue (S156; probability = 0.77; Fig. 5C and D) whose side chain reaches toward the AURKC ATP-binding site (marked by the inhibitor BRD-7880; (Abdul Azeez et al., 2019); Fig. 5C and D). These data suggest that positive selection in the intrinsically disordered region of the ancient protein kinase domain of AURKC regulates error-correction, or other meiotic functions, by modulating kinase activity.

Double RWD domains mediate the formation of CENP-OP heterodimers, allowing recruitment of the CENP-OPQUR complex to centromeres (Pesenti et al., 2018; Schmitzberger and Harrison, 2012). Aligning the FREEDA-annotated CENP-O and -P C-terminal RWD domains to the experimentally solved human CENP-OPQUR complex (Yatskevich et al., 2022) shows that residues under positive selection are on opposite sides of the CENP-OP heterodimer (Fig. 5E) and therefore unlikely to impact heterodimerization. In yeast, C-terminal RWD domains of CENP-O and -P orthologues bind to CENP-Q and -U orthologues to form the COMA complex (Hinshaw and Harrison, 2019; Schmitzberger and Harrison, 2012). We were unable to reliably align mouse CENP-Q and -U to the human CENP-OPQUR complex, likely due to long unstructured regions in CENP-Q and -U, but the striking pattern of residues under positive selection in C-terminal RWD domains facing the outside of the heterodimer suggests adaptive evolution in binding nearby centromeric components. We find that CENP-Q and -U also evolve under positive selection in rodents (Fig. 3), suggesting that positive selection regulates interactions between CENP-OPQUR complex components. Altogether, these analyses of multiple centromere proteins demonstrate how FREEDA-annotated structures can be used to generate hypotheses for how positive selection might regulate essential protein functions.

### Experimental testing of functional protein innovation

To test our hypothesis that intrinsically disordered regions in ancient protein domains regulate their essential functions, we chose to focus on CENP-O because FREEDA shows positive selection flanking α-helices and β-sheets of both rodent and primate C-terminal RWD domains (Fig. S6A-D), and centromere binding provides a straightforward functional assay. We used mouse oocytes for these experiments because they are an established model system for centromere drive, the most likely selective pressure sculpting evolution of centromeric proteins, and thus a natural context to probe for functional protein innovation. To create an evolutionary mismatch (see Introduction), we introduced GFP-tagged full-length mouse (control) or rat (divergent) CENP-O at similar expression levels (Fig. S7A). Mouse CENP-O localized to centromeres as expected, but rat CENP-O was nearly undetectable at mouse centromeres (Fig. 6A and B), indicating functional innovation in centromere binding. To test if the C-terminal RWD domain is responsible for that innovation, we compared three chimeric rat CENP-O constructs with different regions of mouse CENP-O: N-terminal (N-terminal tail and N-terminal helix), central (N-terminal RWD domain and central helix), or C-terminal (C-terminal RWD domain) (Fig. 6A and B; Fig. S7A). Only the mouse C-terminal RWD domain could rescue, albeit not fully, the localization of rat CENP-O to mouse centromeres. In an inverse experiment, a chimera of mouse CENP-O with the rat C-terminal RWD domain failed to localize to mouse centromeres (Fig. 6C and D; Fig. S7B). Together, these results demonstrate that mouse-specific innovation in the C-terminal RWD domain is required for CENP-O binding to mouse centromeres. Within this domain, 10 out of 13 residues that differ between mouse and rat are putatively adaptive (probability ≥ 0.5; Fig. S8). Almost all (9/10) of these residues flank α-helices or β-sheets (+/− 1 amino acid), consistent with our hypothesis that positive selection in intrinsically disordered regions regulates functions of ancient domains. Swapping 5 of the most likely adaptive residues in the mouse C-terminal RWD domain to those found in rat did not reduce centromere localization of mouse CENP-O (Fig. S9A-C), highlighting the difficulty in attributing innovation to specific residues given the number of possible combinations as well as the potential for epistasis (Starr and Thornton, 2016). Altogether, we show that our fully automated molecular evolution pipeline can guide experimental testing of functional protein innovation.

**Figure 6.**
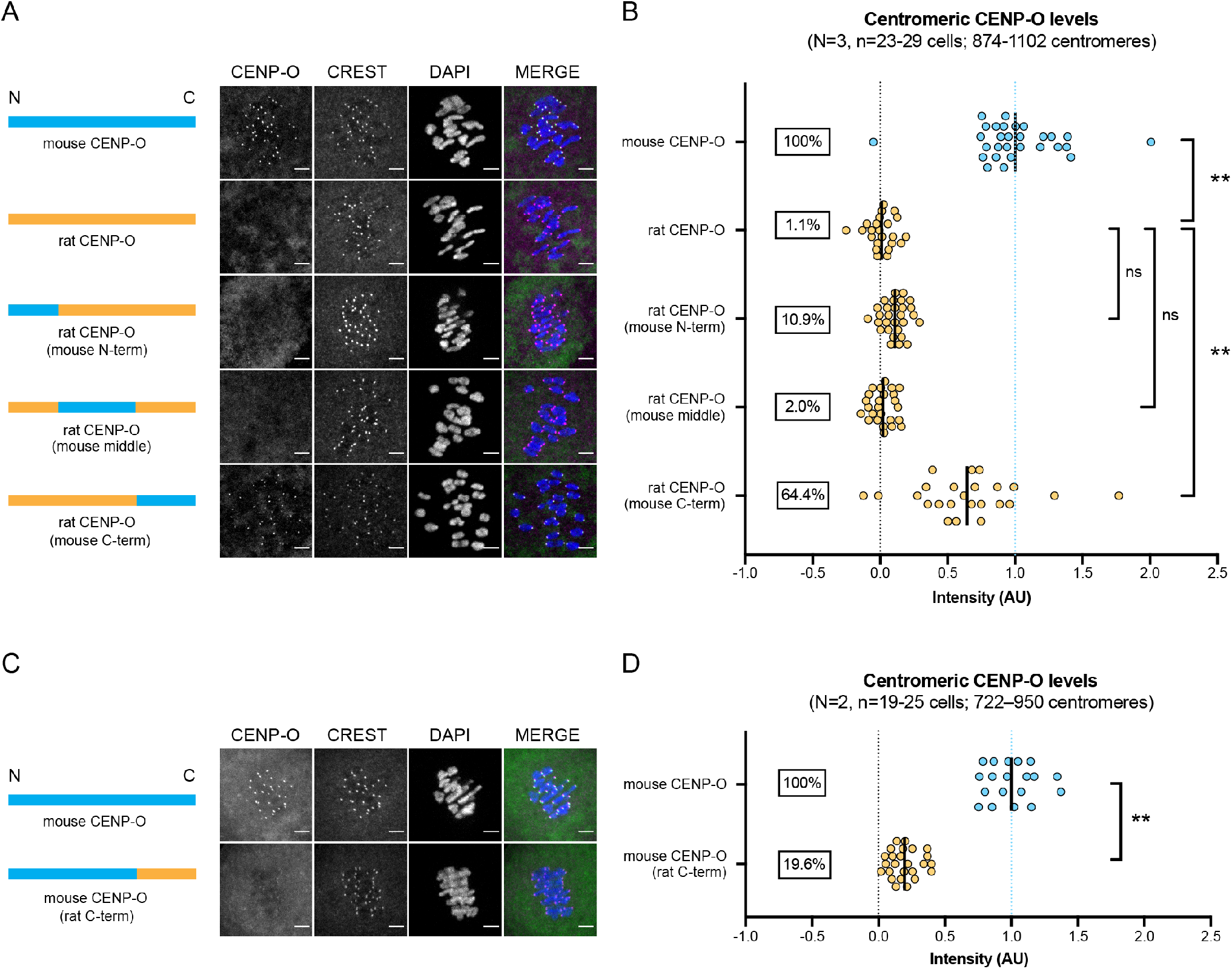
Experimental evidence of functional innovation in the CENP-O C-terminal RWD domain. Mouse oocytes expressing mouse, rat or chimeric CENP-O-GFP were fixed in meiosis I and stained for centromeres (CREST) and DNA (DAPI). Images (**A, C**) show maximum intensity projections; scale bars 5μm. Graphs (**B, D**) show CENP-O-GFP intensity at centromeres; for each construct, n ≥ 722 centromeres from ≥ 19 cells from 3 (B) or 2 (D) independent experiments. Each spot represents one cell; bars: mean intensities; ** p-value < 0.0001, ns: not significant, One-way ANOVA with Tukey’s multiple comparison test (B) or two-tailed student T-test (D).

## Discussion

The motivation to develop FREEDA was to catalyze participation of the cell biology community in testing functional consequences of protein innovation. We demonstrate that detection of positive selection, which implicates functional innovation, can be fully automated by compiling widely used bioinformatic and molecular evolution tools into a single pipeline (Fig. 1). FREEDA’s simple and user-friendly GUI makes it a suitable entry point for experimentalists who may have limited programming skills (Fig. 2). Moreover, by leveraging the ever-growing pool of newly sequenced but not yet annotated genomic assemblies, FREEDA bypasses the requirement for obtaining tissue samples and cloning the genes of interest in order to have sufficient numbers of orthologues from closely related species to detect positive selection. Nevertheless, as with any fully automated tool, FREEDA has limitations. First, by inferring orthologues based on the annotated reference sequence, rather than experimentally validated transcripts, FREEDA does not account for tissue-specific splicing, shifts in intron-exon junctions, or the use of alternative exons. Despite this caveat, using independently annotated rat sequences as reference led to the same result as using mouse annotations in 85/89 centromeric genes (note that relatively poorer rat genome annotation quality prevented reliable input generation for 15 genes; Supplementary Table 3). Nevertheless, isoforms that substantially differ from the reference coding sequence might interfere with accurate detection of positive selection. As an example, annotated variants of the rat NUP37 nucleoporin substantially differ at their C-termini from the reference mouse NUP37 sequence, suggesting the use of alternative exons (Additional Supplementary Materials), which likely led to inconsistent signals of positive selection (p=0.510 with mouse as reference vs. p=0.0499 with rat as reference; Supplementary Table 3). Second, to prioritize computational speed and reduce output complexity, FREEDA does not test for possible recombination events, known to increase the probability of false positives when recombination rates are high (Anisimova et al., 2003). However, estimated recombination frequencies are lower in mammals compared to other animals, yeast and protozoa (Stapley et al., 2017; Wilfert et al., 2007). Third, while FREEDA can robustly resolve gene duplications present in the ancestor of a selected taxon (e.g., primates), caution is advised when analyzing lineage-specific genes. For example, primate *MICA* (MHC class I chain-related gene A) is known to have duplicated from *MICB* in the common ancestor of Hominoids and Old World Monkeys (Florio et al., 2018). Therefore, searching for *MICA* orthologues across the entire primate taxon yields *MICB* coding sequence in New World Monkeys (Additional Supplementary Materials). In case lineage-specificity is suspected, we suggest using the “Exclude species” option (Fig. 2A). Fourth, FREEDA is designed to test for recurring (pervasive) positive selection acting on the entire taxon (e.g., primates) rather than episodic selection that leads to adaptation in a specific branch (e.g., Apes). Finally, although FREEDA can detect residues under positive selection in carnivores (*Carnivora*) and birds (*Phasianidae*), in addition to rodents and primates, mapping them onto protein structures is not yet fully supported.

Our analyses of genes with known evolutionary histories demonstrate that FREEDA is reliable. Building on this validation, we provide the most detailed characterization of positive selection at rodent centromeres to date. Consistent with previous analyses (Kumon et al., 2021), we find pervasive evolutionary innovation in domains of centromeric proteins that do not directly touch DNA, such as RWD (Fig. 3). Therefore, our data support the idea that fitness costs of centromere drive are suppressed by innovation in protein-protein interactions (Kumon et al., 2021; Rosin and Mellone, 2016) as well as protein-DNA interactions (Henikoff et al., 2001; Malik et al., 2002; Vermaak et al., 2002). Furthermore, by mapping regions under positive selection onto protein structures, we derive a hypothesis that positive selection acts on intrinsically disordered regions within ancient domains to impact essential functions. For example, recurrent amino-acid changes within the intrinsically disordered CXXC motifs could regulate MIS18 complex formation. Similarly, recurrent changes in loops of the ancient AURKC protein kinase domain could modulate kinetochore-microtubule attachment dynamics specifically in meiosis I. Both centromere assembly and microtubule detachment (in meiosis I) represent mechanisms potentially hijacked by selfish centromeres (Akera et al., 2019; Henikoff et al., 2001; Rosin and Mellone, 2016; Wu et al., 2018). These analyses provide a starting point for future experiments probing the functional impacts of innovation, including testing whether positive selection reduces fitness costs associated with centromere drive (reviewed in (Dudka and Lampson, 2022)).

Previous experiments in fruit flies, using evolutionarily mismatches of an intrinsically disordered region (L1 loop) within the ancient histone fold domain of Cid^CENP-A^, suggested functional innovation in a DNA-binding domain of a centromere protein (Rosin and Mellone, 2016; Vermaak et al., 2002). Here, we show that positive selection in CENP-O regulates centromere binding via intrinsically disordered regions within an ancient C-terminal RWD domain (Fig. 6), which does not interact with DNA (Pesenti et al., 2022; Yatskevich et al., 2022). While we are unable to pinpoint the exact residues responsible for functional innovation, the observation that most recurrently changing residues are within intrinsically disordered regions of that domain supports our hypothesis. CENP-O is expected to dock the CENP-OPQUR complex to centromeres (Eskat et al., 2012; Pesenti et al., 2018), promoting kinetochore-microtubule attachment stability (Amaro et al., 2010; Bancroft et al., 2015; Chen et al., 2021; Hori et al., 2008; Singh et al., 2021). Therefore, we propose that innovation in the CENP-O C-terminal RWD domain modulates interactions with other centromeric components (possibly CENP-Q and -U) necessary to form a stable CENP-OPQUR complex at centromeres (Foltz et al., 2006; Hori et al., 2008; Kagawa et al., 2014; Minoshima et al., 2005; Okada et al., 2006; Pesenti et al., 2018), potentially stabilizing kinetochore-microtubules to counteract destabilizing activities associated with driving centromeres (Akera et al., 2019; Wu et al., 2018). Future work using centromere drive models (reviewed in (Dudka and Lampson, 2022)) will be needed to experimentally test this idea. Overall, we show how FREEDA can help derive evolutionary hypotheses and guide experimental testing of functional innovation, starting from just a gene name, making it a powerful tool for incorporating evolutionary analyses into cell biology research and generating new insights into essential cellular processes.

## Methods

### Resources and datasets

FREEDA was written in Python and compiled into a stand-alone application using pyinstaller (https://pyinstaller.org/en/stable/). Core packages used for the compilation (Supplementary Table 4) were installed using standard package managers: pip (https://pypi.org/project/pip/) and conda (https://docs.conda.io/en/latest/). All selected genomic assemblies and Ensembl releases used to generate datasets are listed in Supplementary Table 4. All data were collected using a desktop computer iMac (Late 2015) with MacOS Monterey 12.6 (8GB RAM; 4 CPU cores; 2.8 GHz Quad-Core Intel Core i5) or a laptop MacBook Pro (Mid 2014) with MacOS Mojave 10.14.6 (16GB RAM, 2 CPU cores; 3GHz Inter Core i7).

### Input extraction and identification of potential orthologous exons

Both MacOS and Linux releases of FREEDA can be downloaded freely from an open source GitHub repository: https://github.com/DDudka9/freeda/releases. Prior to running the app, MacOS users need to download PyMOL, which renders a three-dimensional result of the FREEDA analysis. To run the pipeline, the user is prompted by the GUI to provide a gene name, select a reference species, and select a location where all the data will be stored. At least 100 GB of storage space is needed to analyze a single taxon (500 GB is recommended). Optionally, the user can: 1) specify the coordinates of residues or domains of interest that will be labeled on the protein structure prediction, 2) customize the BLAST search and orthologue finding (see below), and 3) exclude selected species from the analysis. Advanced user can also specify the codon frequency model used (F3×4 or F3×4 and F61; see PAML manual for details http://abacus.gene.ucl.ac.uk/software/pamlDOC.pdf). The pipeline starts with downloading the reference genome (using NCBI Datasets (Sayers et al., 2021)) and then retrieving all possible UniProt IDs (UniProt, 2021) for a protein encoded by the gene of interest and matching them to AlphaFold database entries (Jumper et al., 2021). Next, FREEDA extracts protein sequence and coding sequence (using pyensembl package https://github.com/openvax/pyensembl/)) from the Ensembl database (Cunningham et al., 2022) and exon sequences and gene sequence (using pybedtools (Dale et al., 2011)) from the downloaded reference genome. Visualization of residues under positive selection requires that the protein sequence of the structural prediction and the protein sequence from the Ensembl database are identical (tested using the Biopython package; (Cock et al., 2009)). If the proteins are not identical, FREEDA performs the analysis without mapping the residues onto structure predictions. The first run triggers downloading of the selected reference genome (e.g. human) and the genomes of closely related species (e.g. *Simiiformes*) and building of local BLAST databases (using BLAST+ applications; (Camacho et al., 2009)). FREEDA queries these databases to find genomic coordinates of putative orthologous regions using tblastn algorithms (default identity threshold is set at 60% but can be increased to 80% by selecting the advanced option “Common domains expected”) and retrieves corresponding nucleotide sequences (using pybedtools) from downloaded related genomes. Overall, these features allow fully automated generation of the input data needed to find orthologous coding sequences in non-annotated genomes.

### Finding orthologous exons

FREEDA performs a multiple sequence alignment of each region found during the BLAST search to the reference coding sequence, BLAST sequences stitched together, the genomic locus these sequences reside in (contig), and the reference gene sequence using MAFFT (Multiple Alignment using Fast Fourier Transform; (Katoh and Standley, 2013). Regions aligning to both the coding sequence and the reference gene sequence are considered as putative exons. To determine if the putative exon is syntenic (resides in a homologous locus), the flanking sequence is compared to the introns of the reference gene separately at 5’ and 3’ ends. An exon is considered syntenic if at least one of the flanking regions is at minimum 75% identical to the reference intron over a stretch of at least 50bp. The identity is calculated as the Hamming distance (the number of different bases in a pair-wise comparison of two aligned sequences; (Hamming, 1950); divided by the sequence length). The putative exon is called as not syntenic and discarded from the analysis if none of the flanking regions reaches the identity threshold and sthe exon itself is less than 80% identical to the reference exon over a stretch of the first 30bp. Since introns are generally less conserved than exons, when the identity of a flanking region is uncertain (66%-75%), a longer sequence is compared.

To increase stringency in detecting synteny and avoid segmental gene duplications, the user can additionally select an advanced option “Duplication expected” that penalizes any exon that is syntenic only at one end (e.g., recent segmental duplication whose introns have not yet diverged significantly). Segmental duplications that lead to duplicated genes residing next to each other (tandem duplications) will not only have similar flanking regions, but might also be difficult to align if residing on the same contig. To ensure robust analysis of tandem duplications, the “Tandem duplication expected” option limits the flanking region of each blast hit (leading to smaller contigs), decreasing the chance of tandemly duplicated genes residing on the same contig. In addition, to avoid retro-duplications (mRNA-derived gene duplications; (Kaessmann et al., 2009)), FREEDA always discards exons that are at least 80% identical to the reference exon but lack the flanking regions (intron-less).

To preserve intron-exon boundaries, each putative exon is given a number and directly aligned to a reference exon of the same number. Therefore, exons do not need to reside on the same contig to form a complete coding sequence, which is helpful when querying genomic assemblies with short contigs. Very small exons (microexons; reviewed in (Ustianenko et al., 2017)) shorter than 18bp cannot be reliably aligned and are discarded from the analysis. If the same putative exon is found on different contigs (e.g., due to duplication), the contig containing fewer putative exons is discarded. If both contigs carry the same number of putative exons (likely due to heterozygosity of the orthologous locus or a very recent duplication), these are compared to corresponding reference exons, and the contig with a higher overall identity of exons is considered orthologous. To allow manual review, all the above-mentioned steps are logged and all the intermediate alignments are saved as raw data (“Raw_data” folder).

### Manual verification of detected orthologues

The ability to distinguish tandem duplication (*H4C1* from *H4C2*) and recent retro-duplication (*KIF4A* from *KIF4B*) was tested by manual BLAST (blastn) of the nucleotide sequence of each orthologue identified by FREEDA (“GENE_raw_nucleotide_alignment.fasta” file) against the primate NCBI gene database (*Simiiformes*; taxid: 314294). For each gene, an orthologous sequence was always the highest scoring hit (by similarity) as opposed to a paralogous sequence. Exceptions were sister species *Aotus nancymaae* and *Callithrix jacchus*, whose identified *KIF4B* coding sequences were more similar to *KIF4A* than *KIF4B*. However, all the *KIF4B* orthologues detected by FREEDA were intron-less, consistent with *KIF4B* being a primate-specific retro-duplication of *KIF4A* (Florio et al., 2018), which FREEDA called correctly. Therefore, we are confident that FREEDA identified *KIF4B* orthologues for all species and not *KIF4A* paralogues (Additional Supplementary Materials).

### Building the multiple sequence alignment and phylogenetic gene tree

Detection of recurring amino acid substitutions requires a gapless, in-frame, multiple sequence alignment. To avoid large gaps (suggesting incomplete coding sequences), FREEDA first removes entire coding sequences that are shorter than 90% compared to the reference sequence. To ensure high quality alignment of the remaining sequences, FREEDA uses a modified MAFFT protocol designed to limit over-aligning errors (Katoh and Standley, 2016). To curate the alignment, FREEDA removes insertions that are defined as regions missing in the reference coding sequence, and deletes stop codons (including premature ones). At this point, coding sequences that are less than 69% identical to the reference sequence are discarded as likely too divergent to produce a reliable alignment (based on (Jeffares et al., 2015; Sievers et al., 2011) or mis-aligned. Additionally, remaining small gaps (deletions) and ambiguously aligned codons are removed from the alignment (using Gblocks; (Castresana, 2000; Talavera and Castresana, 2007)). Finally, to ensure that the curation process did not alter the open reading frame of the aligned sequences, FREEDA compares the identity of the translated reference sequence within the curated alignment to the original reference protein sequence from Ensembl database. Since sequence-wide identity checks explained above might miss small, mis-aligned regions in large genes, FREEDA additionally translates all curated coding sequences and removes those that contain contiguous stretches of a least 10 non-synonymous substitutions when compared to the reference sequence. Final multiple nucleotide sequence alignment is then used to build a phylogenetic gene tree (using RAxML; (Stamatakis, 2014)), which guides the widely used CODEML program from the PAML suite (Phylogenetic Analysis by Maximum Likelihood; (Yang, 2007)) to infer recurrent amino acid substitutions suggestive of positive selection. We urge the user to manually verify the final protein alignment (“GENE_protein_alignment.fasta” file), ensuring that there are no obvious mis-aligned regions before considering the results of the PAML analysis (e.g., using the free software Unipro UGENE; (Okonechnikov et al., 2012)). In case of apparent misalignments, we suggest simply re-running the analysis using the “Exclude species” option (Fig. 2A).

### Detection of positive selection

To detect positive selection, FREEDA relies on the rate ratio of non-synonymous (dN) to synonymous (dS) substitutions (dN/dS > 1 suggests positive selection). However, most genes contain conserved regions that evolve under purifying selection (dN/dS < 1), which usually decreases gene-wide dN/dS below 1. Therefore, to find specific regions under positive selection, FREEDA uses “site models” of the CODEML program that allow for varying dN/dS between different codons. Each model describes a set of parameters (including dN/dS per codon; for details, see the official PAML guide; http://abacus.gene.ucl.ac.uk/software/pamlDOC.pdf or a beginners guide (Jeffares et al., 2015) and either allows for sites (codons) with dN/dS ratio >1 (signature of positive selection; M8 and M2a models) or not (null hypothesis; M7 and M1a models). Using a maximum likelihood approach, CODEML then fits the parameters estimated from the data to each model. Significantly more likely fit (based on the Likelihood Ratio Test, “LRT”) to the model that allows for codons with dN/dS >1 indicates the presence of sites evolving under positive selection. Bayesian statistics (BEB; Bayes Empirical Bayes) are then used to find these specific codons. FREEDA outputs the key results of the CODEML analysis within the GUI’s “Results window” (LRT value for M7 vs M8 comparison, p-value, and number of codons under positive selection). Additionally, the results of the M1a vs M2a comparison and the identity of specific codons under positive selection are saved in an excel sheet (“Results_sheet” folder).

### Visualization of residues under positive selection

FREEDA maps the protein sequence of the reference species from the multiple sequence alignment to the expected reference protein sequence and if appropriate, introduces gaps that represent residues excluded from the analysis (“GENE_protein_alignment.fasta” file). Based on that mapping, FREEDA provides both 2D and 3D visual representations of residues evolving under positive selection. 2D bar graphs are provided in the “Graphs” folder. These graphs display the positions of recurrently changing residues (“Posterior mean omega”, top) and the probability of positive selection acting on each codon (“Prob. positive selection”, middle, probability 0-7-1.0; “High prob. positive selection”, bottom, probability 0.9-1.0). Codons excluded from the analysis are marked in grey. 3D representation of residues under positive selection is found in the “Structures” folder, provided that the prediction model from the AlphaFold database matches the protein sequence extracted from the Ensembl database. For clarity, only residues with high probability (≥ 0.9) of positive selection are mapped, and their side chains are shown. The residues excluded from the analysis are colored in grey, and the N-terminal and C-terminal ends are labeled. Additionally, any domain annotation available in the Interpro database (Blum et al., 2021) is automatically marked with distinct color and labeled allowing quick visual identification.

### Manual alignment of structural prediction models

FREEDA-annotated protein structure prediction models from AlphaFold designated by their UniProt entries (Mm MIS18β - A2AQ14; MmAURKC - O88445; MmINCENP - Q9WU62; MmCENP-O - Q8K015; MmCENP-P - Q9CZ92) were aligned to PDB entries (SpMIS18 – 5HJ0; HsAURKC – 6GR8; HsINCENP – 6GR8; HsCENP-OP – 7PB8) using an aligner module in PyMOL. Briefly, a FREEDA-annotated structure (e.g., “Aurkc_Mm.pse” found in “Results-Current-Date/Results/Structures”) was opened in PyMOL, the selected PDB entry was downloaded (e.g., “fetch 6GR8”), and the two structures were aligned (e.g., “align Aurkc_Mm, 6GR8”). The align module first aligns protein sequences and then superimposes their structures, returning RMSD (Root-Mean-Square-Deviation). Lower RMSD values indicate better alignment. All alignments presented here returned RMSD below 2 Angstroms.

### Generation of CENP-O constructs

All CENP-O coding sequences were cloned from testis or liver samples. The use of rat CENP-O to represent a divergent orthologue was motivated by the availability of a rat (*Rattus norvegicus*) tissue sample for cloning. Chimeric CENP-O constructs were designed based on their three-dimensional structure (AlphaFold database). Tissue was mechanically homogenized and total mRNA was isolated using TRIzol reagent (Invitrogen, 15596026), cDNA was prepared using reverse transcription (SuperScript III First-Strand Synthesis System; 18080051), amplified using construct-specific PCR primers (KAPA HiFi Hot Start plus dNTPs; Roche; KK2502), and inserted into the pGEMHE plasmid backbone (In-fusion kit; Takara; 638948). Each CENP-O construct was tagged with GFP at the C-terminus, separated by a linker of 5 glycines. Site-directed mutagenesis was performed using the Quik-Change Multisite Directed Mutagenesis kit (Agilent; 200515) to introduce point mutations. The identity of all constructs was confirmed using Sanger sequencing of the entire coding sequence, including the reporter gene.

### Oocyte isolation, microinjection and *in vitro* maturation

*Mus musculus* mice (CF-1 strain) were purchased from Envigo NSA stock # 033. Females were primed with 5U of PMSG (Pregnant Mare Somatic Gonadotropin; Calbiochem; 367222) injected into the intra-peritoneal cavity 44-48h prior to oocyte collections to induce superovulation. The ovaries were isolated using M2 medium (Sigma, M7167) supplemented with 2.5 mM of the maturation-blocking phosphodiesterase 3 inhibitor milrinone (2.5mM; Sigma Milipore M4659). GV (Germinal-Vesicle) oocytes were collected, denuded mechanically from cumulus cells, and incubated for at least 1h prior to microinjection on a hot plate (38°C) under mineral oil (FUJIFILM Irvine Scientific; 9305). Oocytes were then microinjected with ~5 pl of mRNAs in M2 medium with 2.5 mM milrinone and 3mg/mL BSA at room temperature with a micromanipulator TransferMan NK 2 (Eppendorf) and picoinjector (Medical Systems Corp.). Oocytes were then incubated in 30-50ul drops of Chatot-Ziomek-Bavister medium (CZB; Thermo Fisher, MR019D) under mineral oil (Sigma Milipore M5310) at 37.8°C and 5% CO2 (Airgas) for 16h to allow protein expression. Concentration of mRNA (15-30ng/μl) used was selected by ensuring similar cytoplasmic expression. mRNAs were synthesized using the T7 mScriptTM Standard mRNA Production System (CELLSCRIPT; C-MSC100625).

### Immunofluorescence imaging

Oocyte maturation was induced *in vitro* by washing out milrinone 7.5 hours before fixation. MI oocytes were fixed in freshly prepared 2% paraformaldehyde in PBS (pH 7.4) with 0.1% Triton X-100 for 20 min at room temperature (RT), permeabilized in PBS with 0.1% Triton X-100 for 15 min at RT, placed in blocking solution (PBS with 0.3% BSA and 0.01% Tween-20) overnight at 4°C, incubated 1h with primary antibody in blocking solution, washed 3 times for 15 min each, incubated 1h with secondary antibody, washed 3 times for 15 min each, and mounted in Vectashield with DAPI (Vector, H-1200) to visualize chromosomes. Centromeres were labeled with CREST (human anti-human Anti-Centromere Antibody, 1:200, Immunovision, HCT-0100) and an Alexa Fluor 594–conjugated goat anti-human secondary antibody (ThermoFisher, A-110014). Confocal images were collected as 31 z-stacks at 0.5 μm intervals to visualize the entire meiotic spindle, using a microscope (DMI4000B; Leica) equipped with a 63x 1.3 NA glycerol-immersion objective lens, an xy piezo Z stage (Applied Scientific Instrumentation), a spinning disk (Yokogawa Corporation of America), and an electron multiplier charge-coupled device camera (ImageEM C9100-13; Hamamatsu Photonics), controlled by MetaMorph software (Molecular Devices). Excitation was done with a Vortran Stradus VersaLase 4 laser module with 405 nm, 488 nm, 561 nm, and 639 nm lasers (Vortran Laser Technology).

### Automated image analysis

Confocal images were analyzed using a custom-built Python-based automated program “Centrocalc” available here: https://github.com/DDudka9/Centrocalc. First, the program performs whole chromosome segmentation using the chromosome channel as a mask by greyscale dilation at a width of 16 pixels to ensure the centromeres are included. Then, a threshold is calculated using the ISODATA method. If no chromosome channel is given, the entire cell is considered. Second, the program identifies centromeres using an approach based on (Vermolen et al., 2008). A difference of Gaussians algorithm is used to isolate spots of 150nm. Third, a local maxima algorithm is used to identify centromeres. Up to 38 spots are chosen (the mouse 2n genome contains 40 chromosomes), separated by a minimum of 2 pixels by Chebyshev distance. The spots must be away from the edges of the image (20 pixels in x and y; 2 pixels in z). Fourth, 3D ellipsoid regions of interest (4×4×3 pixels) are drawn using the local maxima. Background ROIs are drawn as volumes around the centromere ROIs (expanding the 3D ellipsoids by 1 pixel in all dimensions). Overlapping centromere ROIs are resolved by distance, where each pixel is assigned to the closest maxima. Fifth, centromere and background intensities are calculated as average grayscale pixel values and saved. ImageJ ROI files are also created to reference back to the original images. Modifying the Centrocalc source code can customize most of the described parameters.

## Acknowledgements

We would like to thank all the members of the Michael Lampson’s and Mia Levine’s labs (University of Pennsylvania) for scientific discussions, especially Mia Levine who provided critical feedback on the manuscript. We extend special thanks to Piero Lamelza, Hyuk-Joon Jeon and Cara Brand for testing the pipeline. Finally, we would like to acknowledge the vibrant community of the StackOverflow server for providing feedback on the code underlying our pipeline.

## Author Contributions

Conceptualization: D. Dudka and M. A. Lampson, Methodology: D. Dudka and R. B. Akins, Investigation: D. Dudka and R. B. Akins, Writing: D. Dudka and M. A. Lampson, Funding Acquisition: D. Dudka and M. A. Lampson, Resources: M. A. Lampson

