## Supplementary Material for "FREEDA: an automated computational pipeline guides experimental testing of protein innovation by detecting positive selection"

| Feature | Selecton | IDEA | JCoDA | Phyleas Prog | POTION | PosiGene | PoSeiDon | MEGA | DGINN | FREEDA |
|---|---|---|---|---|---|---|---|---|---|---|
| Graphical user interface | YES (Server) | YES | YES | YES (Server) | NO | NO | NO | YES | NO | YES |
| Analysis throughput | candidate based | genome wide | candidate based | candidate or genome wide | genome wide | genome wide | candidate based | candidate or genome wide | candidate based | candidate based |
| Pipeline automation | end-to-end | end-to-end | step-by-step | step-by-step | end-to-end | end-to-end | end-to-end | step-by-step | end-to-end | end-to-end |
| Minimal input required | unaligned coding sequences | aligned coding sequences | unaligned coding sequences | proteins IDs and list of species | unaligned coding sequences | unaligned coding sequences | unaligned coding sequences | unaligned coding sequences | reference coding sequence | gene name |
| Takes advantage of unannotated genomes | NO | NO | NO | NO | NO | NO | NO | YES | YES | YES |
| Finds orthologous sequences | NO | NO | NO | YES | YES | YES | NO | YES | YES | YES |
| Makes multiple sequence alignment | YES | NO | YES | YES | YES | YES | YES | YES | YES | YES |
| Builds phylogenetic tree | YES | YES | YES | YES | YES | YES | YES | YES | YES | YES |
| Detects positive selection at specific sites | YES | YES | YES | YES | YES | YES | YES | YES | YES | YES |
| Visualizes positively selected residues in sequence | YES | YES | YES | YES | YES | YES | YES | YES | NO | YES |
| Visualizes positively selected residues in structure | YES | NO | NO | YES | NO | NO | NO | NO | NO | YES |

**Supplementary Figure 1. Features of published automated pipelines used to detect positive selection**.

Only automated pipelines are represented: Selecton (Stern et al., 2007), IDEA (Egan et al., 2008), JCoDA (Steinway et al., 2010), PhyleasProg (Busset et al., 2011), POTION (Hongo et al., 2015), PosiGene (Sahm et al., 2017), PoSeiDon (Holzer and Marz, 2021), MEGA (Tamura et al., 2021), DGINN (Picard et al., 2020).

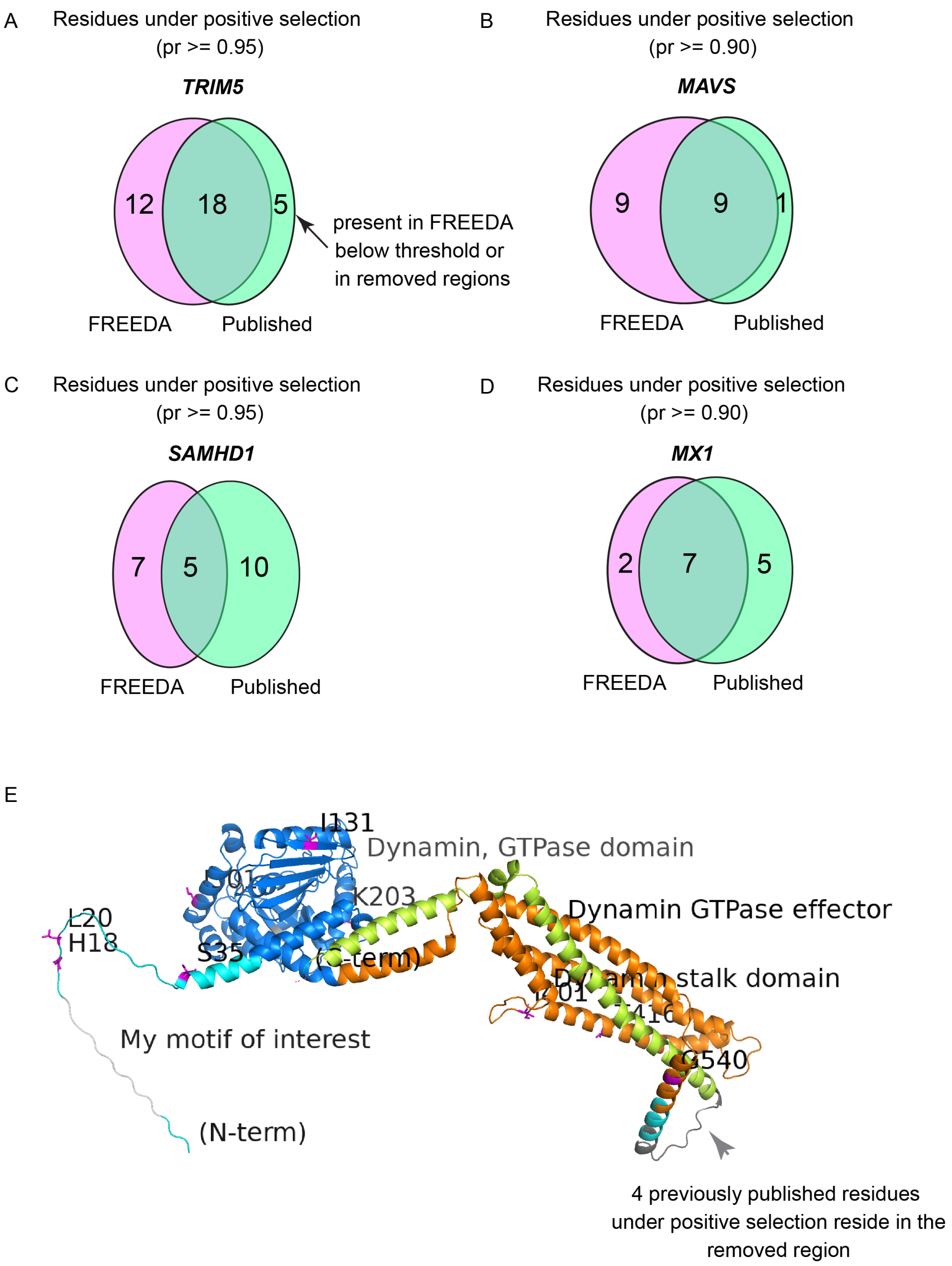


**Supplementary Figure 2. Pipeline validation at the level of single residues**
**A**) *TRIM5* analysis of 16 species with 96% CDS coverage as compared to (Sawyer et al., 2005; van der Lee et al., 2017). **B**) *MAVS* analysis of 19 species with 97% CDS coverage as compared to (Patel et al., 2012). **C**) *SAMHD1* analysis of 18 species with 92% CDS coverage as compared to (Laguette et al., 2012; Lim et al., 2012; van der Lee et al., 2017). **D**) *MX1* analysis of 18 species with 97% CDS coverage as compared to (Mitchell et al., 2012). See Supplementary Table 2 for detailed analyses. Only number of residues from M8 vs M7 analysis are reported (see Methods for details). Comparisons are made using similar phylogenetic branches of *Simiiformes* (Hominoids, Old World Monkeys and New World Monkeys) in most cases. Probability thresholds are set to match those used by the referenced studies. Number of species used varies due to elimination of incomplete sequences from FREEDA analyses. Magenta elipses - number of sites matching the threshold found by FREEDA; green elipses - number of published sites. E) Raw FREEDA-annotated image of the MxA protein. Note the region removed from the analysis due to alignment uncertainty (dark grey arrowhead) where 4 residues under positive selection have previously been identified (Mitchell et al., 2012).

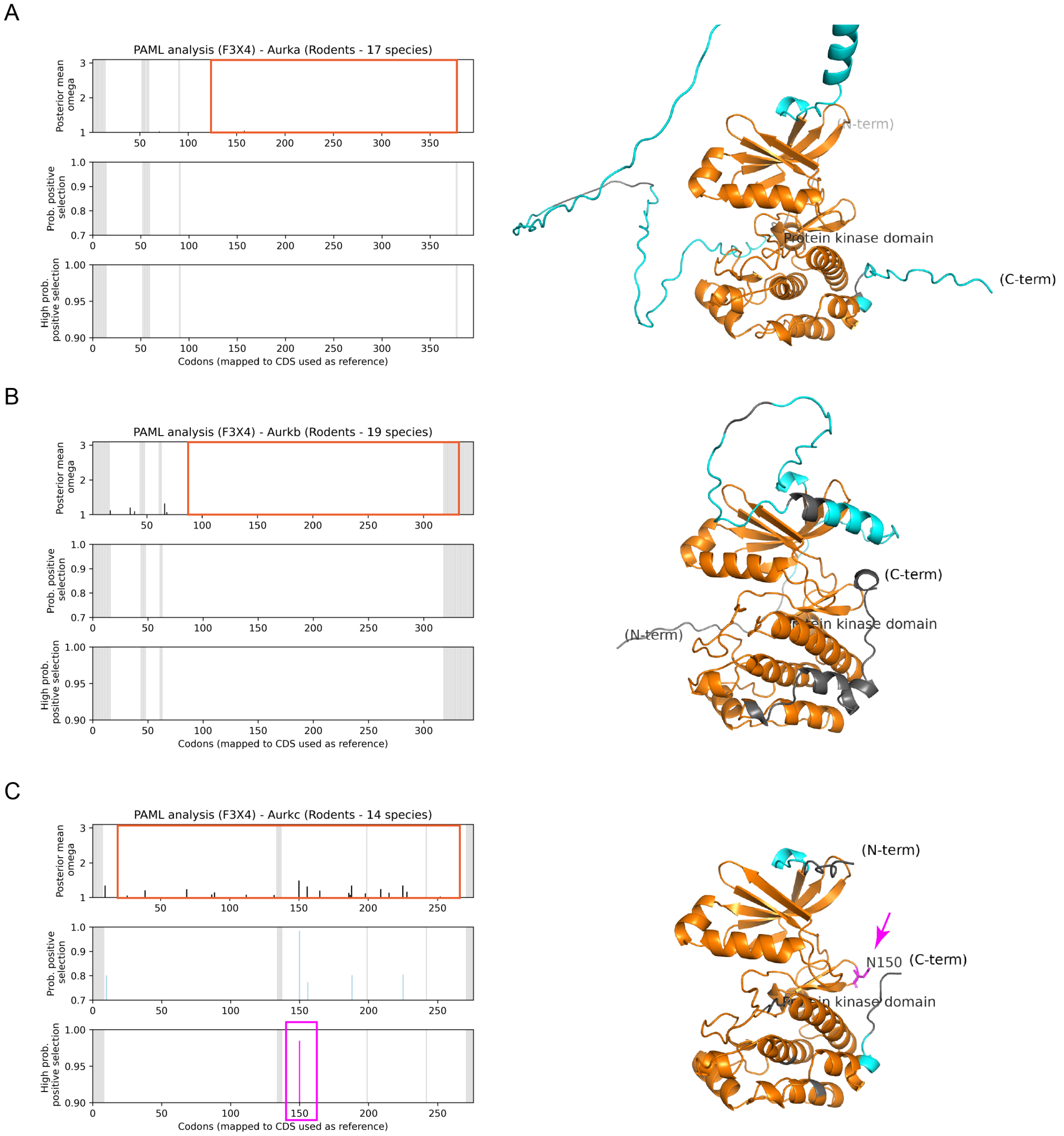


**Supplementary Figure 3. Aurora kinases differ in the number of recurrently changing residues**
Recurrently changing residues, mapped onto reference coding sequences, are not detected in AURKA (**A**) and AURKB (**B**), as compared to AURKC (**C**). Orange frames: protein kinase domain. Structural models highlight high structural conservation of Aurora protein kinase domains (orange). Parts of the AURKA N-terminus were cropped to allow better visualization and comparison of kinase domains.

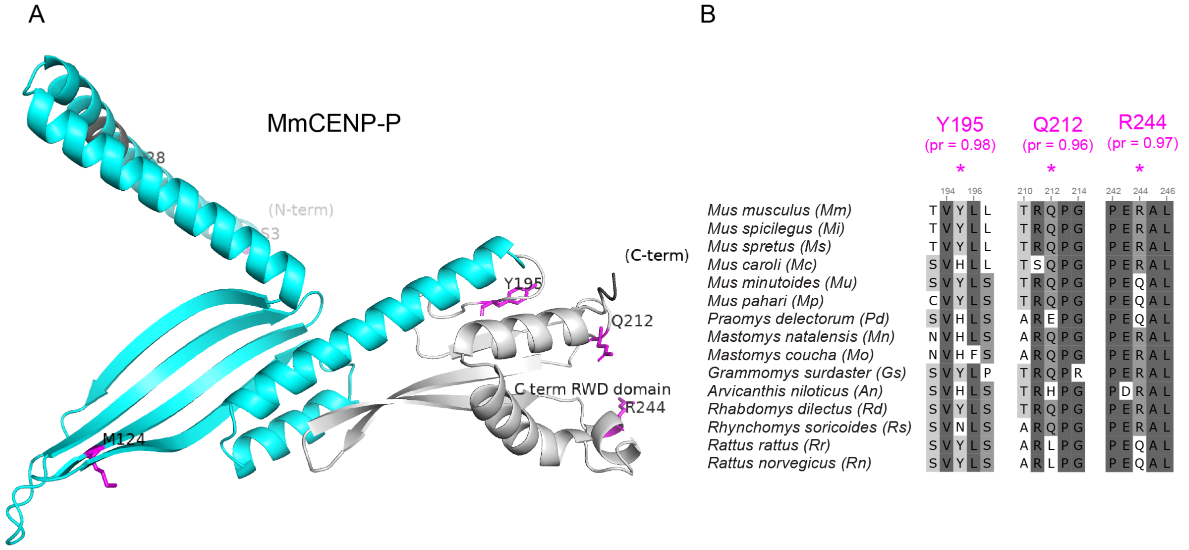


**Supplementary Figure 4. Positive selection in the CENP-P C-terminal RWD domain**

**A**) Annotated structural prediction model of MmCENP-P, generated automatically by FREEDA and visualized in PyMOL without manual modifications. The C-terminal RWD domain (indicated by the user within the GUI) is in grey. (**B**) Three residues (magenta) with high probability of positive selection (≥ 0.9) are shown in snippets of the multiple sequence alignment in *Murinae*. Dark grey: highly conserved residues, grey: less conserved residues, white: non-synonymous substitutions.

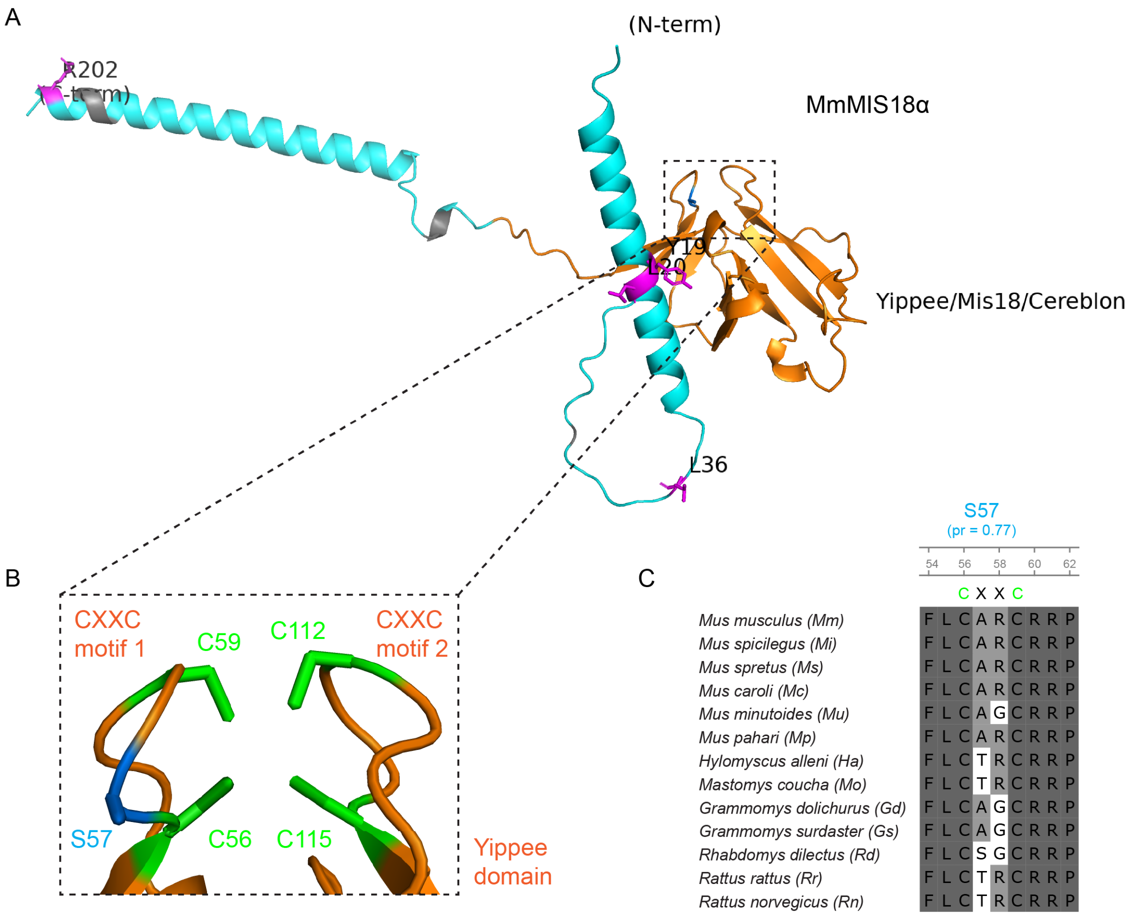


**Supplementary Figure 5. Positive selection in the CXXC motif of the Yippee domain of MIS18α**

**A**) Annotated structural prediction model of mouse MIS18α (MmMIS18α), generated automatically by FREEDA and visualized in PyMOL without any manual modifications. Orange: Yippee domain, magenta: highly likely adaptive residues (probability $\geq$ 0.9). **B**) Enlarged CXXC motifs with the S57 residue (blue) within motif 1. Green: conserved cysteine residues. **C**) Snippet of the multiple sequence alignment of CXXC motif 1 in *Murinae*. Dark grey: highly conserved residues, grey: less conserved residues, white: non-synonymous substitutions.

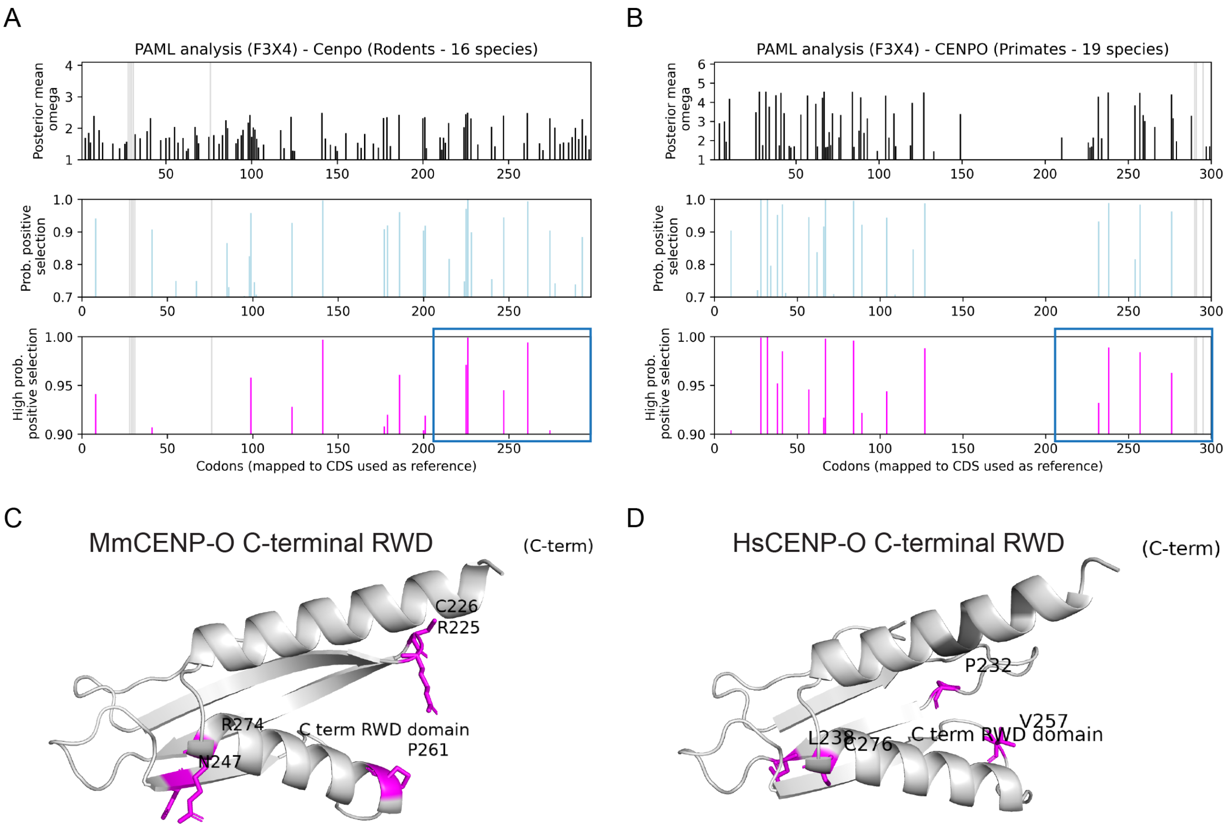


**Supplementary Figure 6. Recurrent innovation in intrinsically disordered regions of rodent and primate CENP-O C-terminal RWD domains**

Recurrently changing residues in rodent **(A)** or primate **(B)** CENP-O mapped to the mouse or human CENP-O coding sequence, respectively, by FREEDA. Blue frames were manually added to outline C-terminal RWD domains. **C** and **D**) FREEDA-annotated C-terminal RWD domains of mouse (C) and human (D) CENP-O. Magenta: highly likely adaptive residues (probability ≥ 0.9).

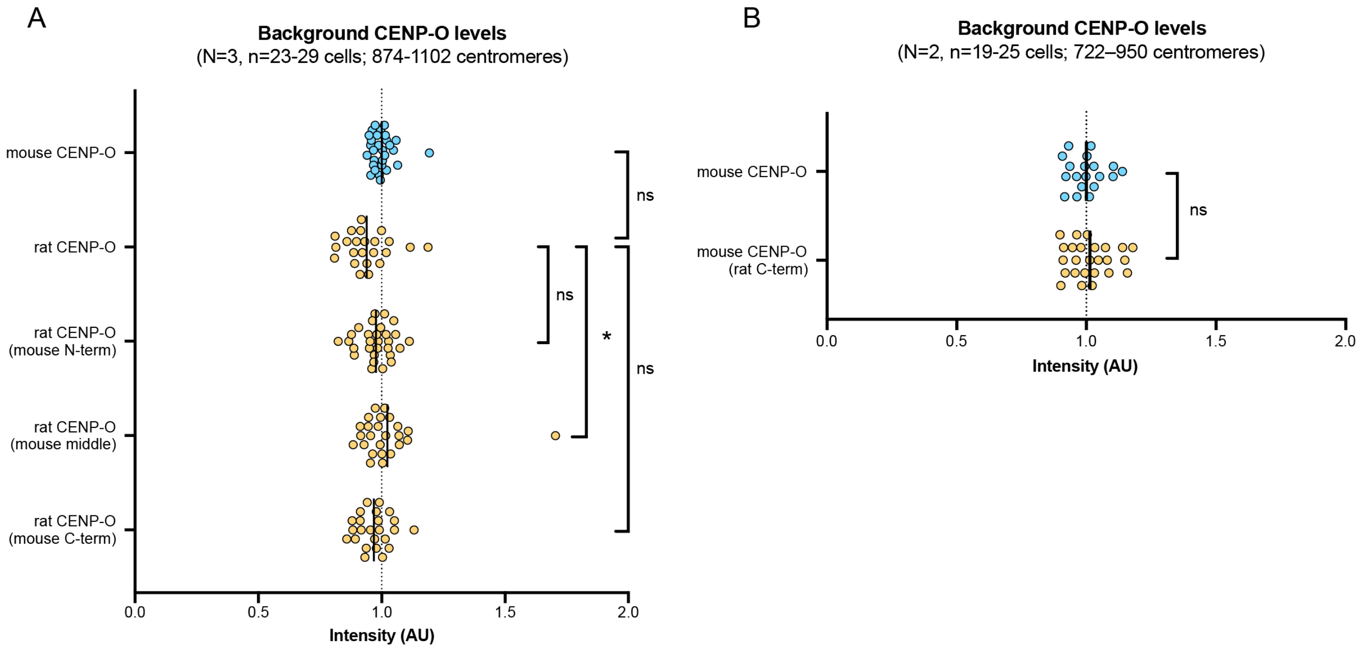


**Supplementary Figure 7. Expression levels of microinjected CENP-O-GFP are equal between constructs**

Quantification of CENP-O-GFP levels in cytoplasm for constructs analyzed in Figure 6A and B **(A)** and Figure 6C and D **(B)**. Each spot represents one cell. For each construct, 38 cytoplasmic ROIs per cell were analyzed (see Methods for details) from ≥ 19 cells from 3 (A) or 2 (A) independent experiments. Bars: mean intensity. * p-value < 0.05; ns: not significant; One-way ANOVA with Tukey's multiple comparison test (A) or two-tailed student T-test (B).

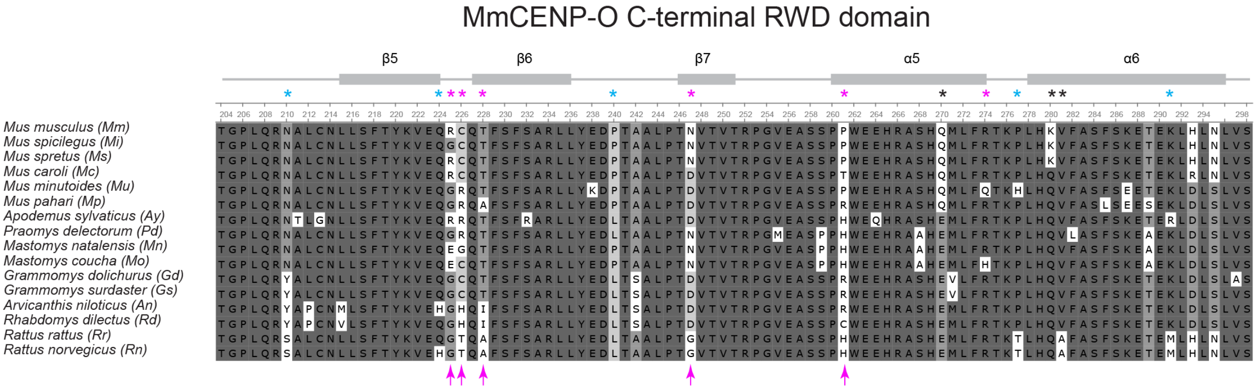


**Supplementary Figure 8. Multiple sequence alignment of the rodent CENP-O C-terminal domain**
Schematic shows α-helices and β-sheets (grey boxes) and intrinsically disordered regions (loops and β-turns: grey lines). Asterisks indicate: highly likely adaptive residues (magenta, probability ≥ 0.9), less likely adaptive residues (blue, probability ≥ 0.5), and residues that differ between mouse and rat but do not evolve under positive selection (black). Magenta arrows indicate residues swapped in the experiment. Alignment: dark grey: highly conserved residues, grey: less conserved residues, white: non-synonymous substitutions.

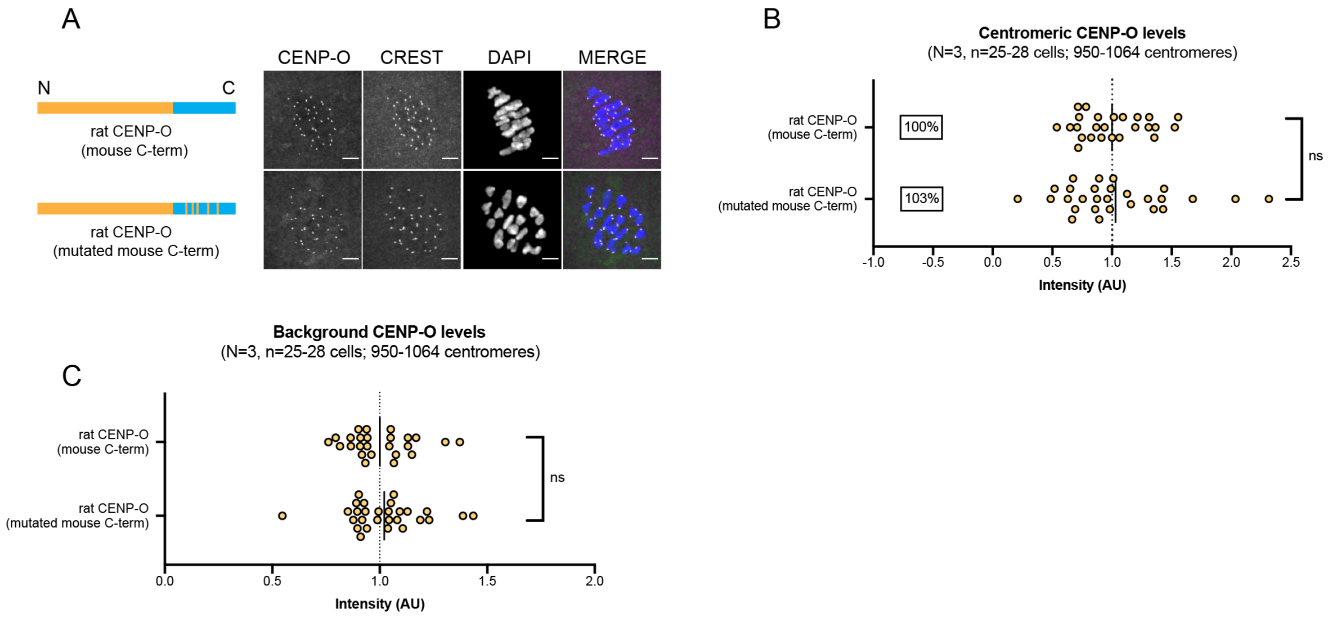


**Supplementary Figure 9. Swapping five residues under positive selection in the CENP-O C-terminal RWD domain is insufficient to reduce centromere binding**

Mouse oocytes expressing the indicated CENP-O-GFP constructs were fixed in meiosis I and stained for centromeres (CREST) and DNA (DAPI). One construct is rat CENP-O with the mouse C-terminal RWD domain, and the other is identical except for 5 mutations in the RWD domain swapping mouse-specific residues under positive selection (probability $\geq$ 0.9) to the corresponding rat-specific residues. Images (**A**) show maximum projections; scale bars 5µm. Graphs show CENP-O-GFP intensity at centromeres (**B**) or in the cytoplasm (**C**); for each construct, n $\geq$ 950 centromeres from $\geq$ 25 cells from 3 independent experiments. Each spot represents one cell; bars, mean intensities; ns: not significant, two-tailed student T-test.

**Supplementary Table 1. FREEDA results analyzing test datasets comprising 23 primate proteins.** Red – genes previously reported to have evolved under positive selection in primates. Green – genes with variable reports of their adaptive evolution. Blue – genes previously reported to not have evolved under positive selection. Related to Fig. S2.

**Supplementary Table 2. Comparison of specific sites under positive selection published previously and those detected by FREEDA in selected genes.** Probabilities of adaptive evolution are provided for each site in both F3X4 and F61 codon frequencies. Related to Fig. S2.

**Supplementary Table 3. FREEDA results analyzing 104 centromeric proteins in rodents.** Results are shown separately using mouse or rat as reference species. Red indicates genes evolving under positive selection. Red font marks genes with inconsistent evidence of positive selection depending on whether mouse or rat is used as reference species. Grey indicates genes that were not analyzed due to poor annotation in the rat genomic assembly. Related to Fig. 3.

**Supplementary Table 4.** List of genomic assemblies and core packages used by FREEDA to detect positive selection.

### Additional Supplementary Materials

FREEDA results for all proteins analyzed in the manuscript are available here: https://drive.google.com/drive/folders/1lN8clIunaPpbs40MRJDRMokSMAMxFkv-?usp=sharing. These include KIF4A, KIF4B, histone H4, MICA, MICB, NUP73 and HERC5 as mentioned in the text. Manually aligned structural prediction models for MIS18A, MIS18B, AURKC, CENP-O and CENP-P are also included.
